# Spontaneous neurotransmission at evocable synapses predicts their responsiveness to action potentials

**DOI:** 10.1101/2020.08.28.271742

**Authors:** Andreas T. Grasskamp, Meida Jusyte, Anthony W. McCarthy, Torsten W.B. Götz, Susanne Ditlevsen, Alexander M. Walter

## Abstract

Synaptic transmission relies on presynaptic neurotransmitter (NT) release from synaptic vesicles (SVs) and on NT detection by postsynaptic receptors. Transmission exists in two principal modes: action-potential (AP) evoked and AP-independent, “spontaneous” transmission. AP-evoked neurotransmission is considered the primary mode of inter-neural communication, whereas spontaneous transmission is required for neuronal development, homeostasis, and plasticity. While some synapses appear dedicated to spontaneous transmission only, all AP-responsive synapses also engage spontaneously, but whether this encodes functional information regarding their excitability is unknown. Here we report on functional interdependence of both transmission modes at individual synaptic contacts of *Drosophila* larval neuromuscular junctions (NMJs) which were identified by the presynaptic scaffolding protein Bruchpilot (BRP). Consistent with its role in organizing the AP-dependent release machinery (voltage-dependent Ca^2+^ channels and SV fusion machinery), most active BRP-positive synapses (>85%) responded to APs. At these synapses, the level of spontaneous activity was a precise predictor for their responsiveness to AP-stimulation. Both transmission modes engaged an overlapping pool of SVs and NT-receptors, and both were affected by the non-specific Ca^2+^ channel blocker cadmium. Thus, by using overlapping machinery, spontaneous transmission is a continuous, stimulus independent predictor for the AP-responsiveness of individual synapses.

## Introduction

Synaptic transmission relies on quantal neurotransmitter (NT) release from synaptic vesicles (SVs) fusing with the plasma membrane at presynaptic active zones (AZs) and on subsequent NT detection by postsynaptic receptors (Südhof, 2013). Transmission is evoked by action-potentials (APs) but can also occur “spontaneously” in the absence of a stimulus (Fatt and Katz, 1952; Kaeser and Regehr, 2014; Schneggenburger and Rosenmund, 2015).

In evoked transmission, APs depolarize the membrane potential, inducing voltage-gated Ca^2+^ channel opening and Ca^2+^-influx. Subsequently, Ca^2+^ binds to an SV-associated Ca^2+^ sensor (synaptotagmin-1 or -2) and induces SV fusion (Littleton and Bellen, 1995; Kochubey et al., 2011; Südhof, 2012a) which depends on the formation of the neuronal SNARE complex consisting of vesicular VAMP2/Synaptobrevin and the plasma membrane proteins SNAP-25 and syntaxin-1 (Südhof and Rothman, 2009; Jahn and Fasshauer, 2012; Rizo, 2018). AP-evoked NT release further requires the SNARE-binding proteins (M)Unc13 and (M)Unc18 (Brose et al., 2000; Richmond and Broadie, 2002; Toonen and Verhage, 2007; Dittman, 2019). Additionally, AZ cytomatrix proteins like Rab3-interacting molecule (RIM), RIM-binding protein (RBP), and ELKS/Bruchpilot (BRP) contribute to AP-evoked transmission by organizing this release machinery (Südhof, 2012b; Held and Kaeser, 2018; Walter et al., 2018).

In various synapses and model systems, spontaneous transmission was shown to utilize SVs, NT receptors, Ca^2+^ sensors and SNARE proteins separate from ones used during AP-evoked transmission (Broadie et al., 1995; Deitcher et al., 1998; Koenig and Ikeda, 1999; Schoch et al., 2001; Sara et al., 2005; Atasoy et al., 2008; Fredj and Burrone, 2009; Groffen et al., 2010; Sara et al., 2011; Ramirez et al., 2012; Bal et al., 2013; Peled et al., 2014; Courtney et al., 2018). Live imaging experiments furthermore revealed spatial segregation of both transmission modes with no or negative correlation between them (Melom et al., 2013; Peled et al., 2014; Reese and Kavalali, 2016; Farsi et al., 2021). Accordingly, spontaneous transmission was suggested to form a communication channel distinct from AP-evoked activity which may fulfil essential roles in synapse development (Kavalali, 2015). However, at the same time, spontaneous transmission also occurs at all AP-evocable synapses where the same pool of SVs was implicated to cycle spontaneously and during AP-stimulation (Groemer and Klingauf, 2007; Hua et al., 2010; Wilhelm et al., 2010). To what extend spontaneous signals in this “mixed channel” carry information of physiological relevance is unknown.

To investigate this “mixed channel” we here focused on the analysis of spontaneous transmission at AP-responsive AZs of the *Drosophila* neuromuscular junction (NMJ) identified by the scaffolding protein BRP. We show that at these AZs, AP-evoked and spontaneous transmission are highly inter-dependent, and the spontaneous activity predicts AP-responsiveness. Both transmission modes depend on an overlapping machinery on both pre- and postsynaptic sides. Based on these data, we propose that spontaneous activity at AP-evocable synapses serves as a scalable, highly uniform readout of connectivity strength likely relevant for synapse maintenance and plasticity.

## Results

One of the difficulties in assessing the relation between spontaneous and AP-evoked transmission has been the inability to simultaneously monitor both activity modes at the level of individual synaptic connections. This was improved by the development of elegant live-imaging approaches to track synaptic activity with high spatial resolution (Peled and Isacoff, 2011; Melom et al., 2013; Peled et al., 2014; Reese and Kavalali, 2016; Tang et al., 2016; Sakamoto et al., 2018; Farsi et al., 2021). We here use such an assay to monitor synaptic activity at the *Drosophila* 3^rd^ instar larval NMJs (type Ib boutons of muscle 4). In this assay, we postsynaptically express the fluorescent Ca^2+^-reporter GCaMP5 which reports on changes in Ca^2+^-levels induced during synaptic transmission due to the Ca^2+^ permeability of open postsynaptic glutamate receptors (Melom et al., 2013; Reddy-Alla et al., 2017) (**Figure 1A,B**). We confirmed that such fluorescent signals reliably report on synaptic activity in combined electrophysiological experiments (**Figure 1 - figure supplement 1A-F**).

**Figure 1.**
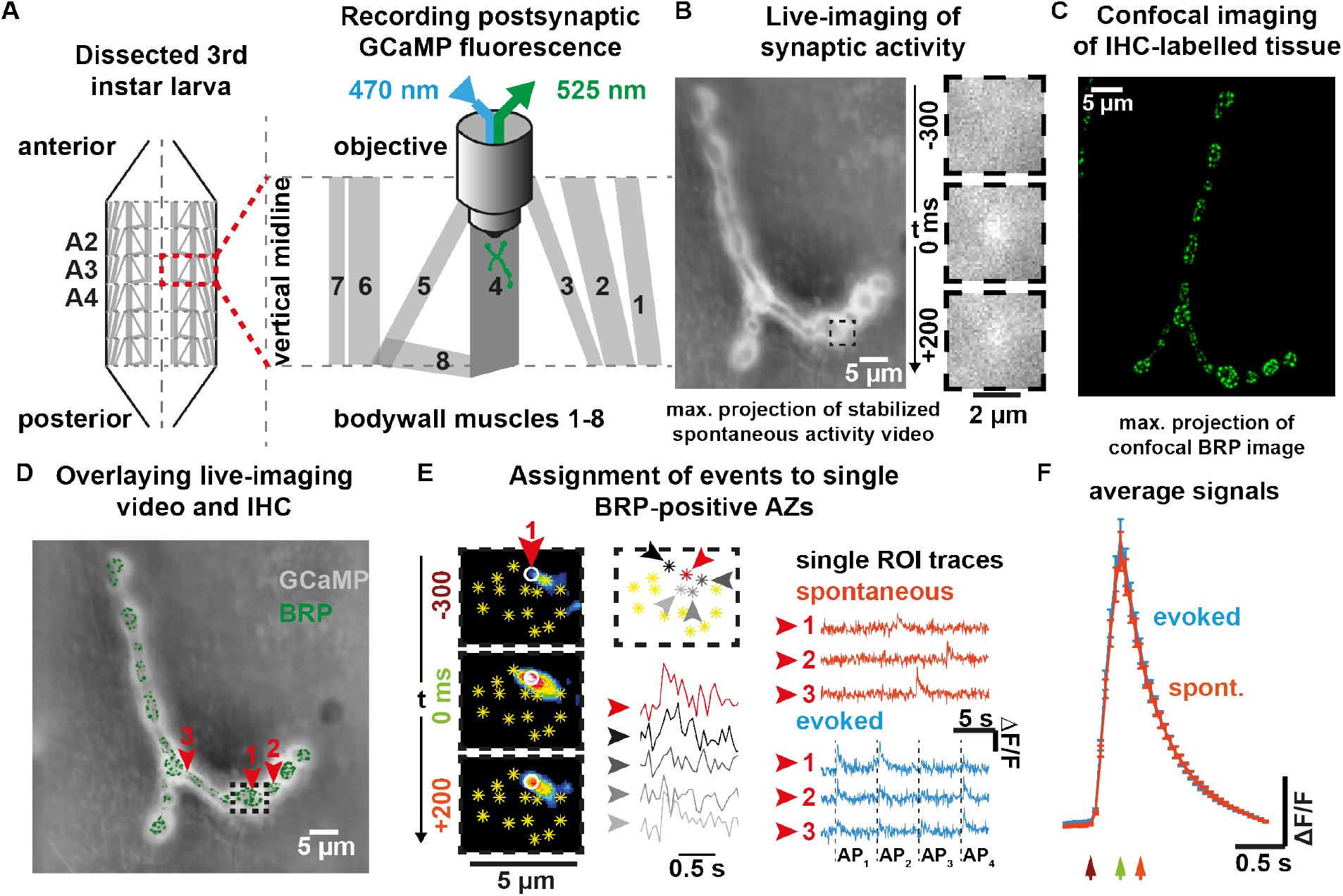
Postsynaptic fluorescence measurements of presynaptic activity with single AZ precision. (**A**) Scheme of experiment to measure Ca^2+^ induced fluorescence signals in *Drosophila* larvae postsynaptically expressing GCaMP. (**B**) Left: Contrast adjusted maximal projection of stabilized 100 s spontaneous activity video; Right: 500 ms time sequence of single event observed in indicated sub-frame. (**C**) Confocal image obtained after immunohistochemistry (anti-BRP staining). (**D**) Maximal projection of 100 s GCaMP video (grey) overlaid with registered confocal BRP image (green). Red arrows indicate locations from which activity shown in (**E**) is read out. (**E**) Detail from black dashed rectangle in (**D**): Left: 500 ms sequence (16-color LUT, contrast adjusted) of a single spontaneous SV fusion event attributed to a single AZ (white circle). Yellow asterisks indicate AZ locations identified by BRP staining & confocal image. Center: event assignment by signal strength within the ROIs. GCAmp trace of the assigned AZs together with the signal at four neighboring AZs are shown. Right: Fluorescence traces from individual AZs in (**D/E**) during either spontaneous (orange) or AP-evoked (blue) recordings. (**F**) Cell-wise mean±SEM (N = 15 animals) fluorescence traces of 670 spontaneous and 2849 AP-evoked events. Arrows point to corresponding times indicated in (E) (dark red: 300 ms before peak, green: at the time of the peak, orange: 200 ms after the peak fluorescence). See also **Figure 1 - figure supplement 1&2**.

We wanted to track transmission at AP-responsive synapses and therefore restricted our analysis to AZs containing the cytomatrix protein BRP which organizes the AP-sensitive release machinery (e.g. voltage gated Ca^2+^ channels and Unc13A release sites (Kittel et al., 2006; Fouquet et al., 2009; Böhme et al., 2016)). To do so, GCaMP movies of live activity were aligned to confocal images obtained post-hoc of the same NMJs stained against BPR (see methods for details) (**Figure 1C,D**). The local temporal GCaMP fluorescence profiles were then read out at equally sized regions of interest (ROIs) placed on all BRP-positive synaptic contacts (**Figure 1E**). Even though postsynaptic Ca^2+^ dispersion caused elevated GCaMP signals at multiple ROIs in response to single events, the signal intensity steeply decreased from the origin to the periphery, allowing signal assignment (highest peak) of temporally scattered spontaneous events to one AZ (**Figure 1E**). Additionally, for AP-evoked responses a distance threshold (2.5 µm) between locations was imposed to prevent that the same event was counted more than once. In principle this could cause a slight under-estimation of AP-evoked activity, but such cases are rare, due to the overall low activity of individual AZs (Muhammad et al., 2015). We first assessed spontaneous activity by imaging signals for 100 s without stimulation. The AP-evoked activity of the same AZs was assessed afterwards in a separate recording where 36 AP stimuli were administered at a frequency of 0.2 Hz (**Figure 1E,F** and **Figure 1 – figure supplement 1G**). The time course and amplitude of NT-induced GCaMP signals were indistinguishable for spontaneous- and AP-evoked release (**Figure 1F**) and GCaMP amplitudes were non-saturating at a Ca^2+^ concentration of 1.5 mM in the external medium ([Ca^2+^]_ext_) (**Figure 1 – figure supplement 1H**), confirming the quantal resolution of this assay (see methods and **Figure 1 – figure supplement 1&-2**).

We first validated to which extent BRP serves as a reliable marker for the AP-responsive AZs we sought to investigate. Using the approach above, local activity maps for either transmission mode at BRP-positive AZs were generated for individual NMJs (**Figure 2A**). While some synapses only showed activity during either the spontaneous or AP-evoked recoding episode, others contributed to both modes (“mixed” AZs) (**Figure 2 – figure supplement 3B)**. Merely observing one type of activity during our recording does not preclude that these AZs might eventually also engage in the other mode at later times or with additional stimuli. Indeed, longer experiments increase the proportion of “mixed” AZs (Melom et al., 2013). However, it is unclear whether eventually (for infinite recording time) all BRP-positive AZs would be expected to engage in both transmission modes or whether some might be dedicated to spontaneous transmission only. To investigate this, we used a survival analysis to extrapolate whether all BRP-positive AZs observed to engage in spontaneous activity during the first recording episode would at some point be expected to additionally be AP-responsive (considered as “death” of a dedicated spontaneous synapse) or whether some AZs might never respond to APs (considered “immortal” for infinite stimuli) (see methods for details). This analysis indicated that even though the fraction of “spontaneous only” synapses decreased exponentially with time, our data was most consistent with a plateau of 21% of AZs predicted to never respond to APs (“immortal” spontaneous only AZ) (**Figure 2 – figure supplement 1**). This constitutes ∼14% of all AZs active throughout the experiment, similar to an estimate of a classic study at the frog NMJ (Zefirov et al., 1995). Thus, while a small fraction of BRP-positive AZs appears dedicated to spontaneous transmission, the vast majority (>85%) of active BRP-positive AZs are AP-responsive (as expected from its role in organizing the AP-dependent release machinery). Analyzing BRP-positive AZs therefore enables us to study the role of spontaneous transmission at predominantly AP-responsive synapses.

**Figure 2.**
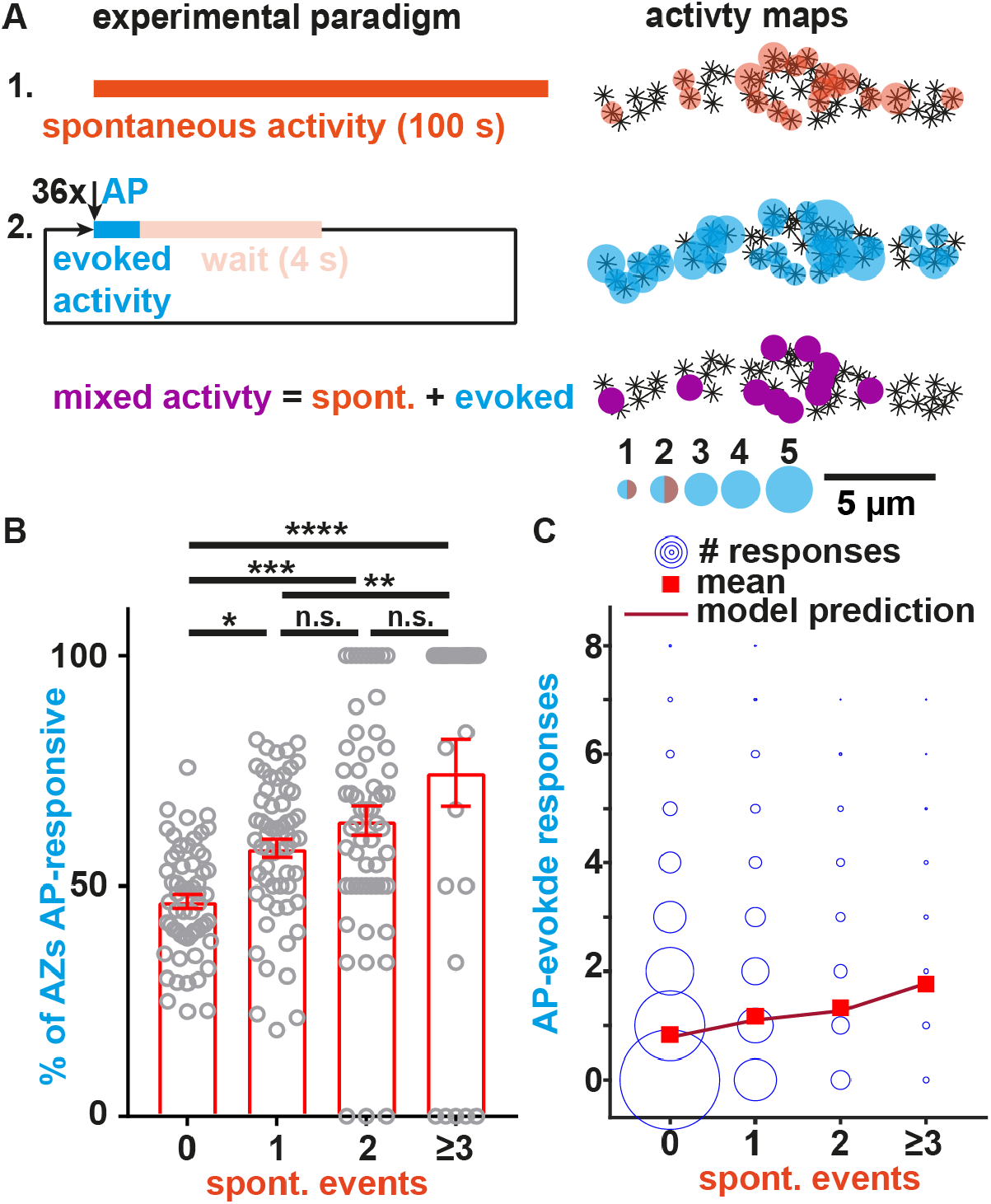
Spontaneous transmission at individual BRP-positive AZs predicts their AP-responsiveness. (**A**) Left: Experimental paradigm. Spontaneous events (top, orange) are recorded for 100 s in isolation before 36 APs are elicited at 0.2 Hz (middle, blue). AZs engaging in both modes during the recording show mixed activity (purple, bottom). Right: AZs of this NMJ are indicated as asterisks and were identified by anti-BRP immunostaining. The local activity profile is indicated by colored circles whose sizes represent the number of spontaneous (orange) or AP-evoked (blue) events (see legend). AZs with mixed activity are shown in purple. (**B**) Animal-wise (N=59 in total) comparison of the fraction of BRP-positive AZs responding to AP-stimulation shown for different numbers of spontaneous events detected at these AZs. Because of the overall low per-AZ activity, fewer animals are found in the groups with more spontaneous events (N(0)=59, N(1)=59, N(2)=58, N(≥3)=29, the group “≥3” pools the responses from AZs with 3 and 4 spontaneous events as only three animals had each one AZ with four spontaneous events). (**C**) Analysis of the relation between the observed AP-evoked and spontaneous transmission events at individual BRP-positive AZs. Events from all imaged AZs in the 59 animals are pooled. The size of the circles relates to the number of observations (between 1 and 3966). Red squares indicate mean number of evoked responses, the line indicates the model prediction (see methods). Number of BRP-positive AZs investigated: n(AZs)=9677, number of animals (N) as in (B). Bars in panel B indicate mean, error bars SEM. * <0.05; ** p<0.01; ***p<0.001; ****p<0.0001; n.s.: not significant. One-way ANOVA with post hoc Tukey’s multiple comparisons test. See also **Figure 2 – figure supplement 1-3**.

We next investigated the relation between both transmission modes at the BRP-positive AZs. We first tested to what extent spontaneous activity predicted the probability of AZs to respond to APs. For this we plotted the fraction of AZs that showed activity at least once during the 36 AP stimuli as a function of the spontaneous activity measured at those AZs prior to stimulation for each animal (59 animals were investigated). This revealed that AZs with (more) spontaneous activity were more likely to respond to APs (Fig. 2B).

We then went on to investigate whether the number of spontaneous events detected prior to stimulation predicted the number of AP-induced responses at individual, BRP-positive AZs. For this we evaluated the experimental data with a generalized linear model using a negative binomial response distribution with a log link. The number of evoked events was modelled a function of spontaneous events and a variable intercept was allowed for each animal (with a random effect). The effect of the number of spontaneous events was strongly statistically significant (p < 0.0001), demonstrating that AP-evoked transmission strongly depended on the spontaneous activity. We furthermore found that a higher spontaneous activity at a given AZ predicted more AP-evoked events (a significant monotonic trend was confirmed using a nonparametric Mann-Kendal trend test, p = 0.045, for details see methods). Importantly, the effect of the number of spontaneous events was also strongly statistically significant (p < 0.0001) with a positive trend (p = 0.045) in separate experiments in which we first measured AP-evoked activity and afterwards measured spontaneous activity (inverse sequence; **Figure 2 – figure supplement 3A,B**), ruling out effects of e.g. activity run-down and fluorescence bleaching. Thus, the spontaneous activity of evocable synapses at rest is a highly reliable predictor of their responsiveness to AP-stimulation.

While above analysis demonstrated that AP-evoked responses of single SZs depended on their spontaneous activity, we next wondered whether -inversely-AP-evoked transmission affected spontaneous activity. To investigate this, we quantified the spontaneous activity “interleaved” between AP-stimuli (**Figure 2 – figure supplement 3A**). While interleaved spontaneous events had similar amplitudes as ones recorded separately, their frequency was strongly reduced (frequencies – seq: 0.0037 Hz/AZ; int: 0.0018 Hz/AZ; N = 9 animals, p = 0.0003; amplitudes – seq: 786.8 a.u.; int: 845.5 a.u.; N = 9 animals, p = 0.13; paired parametric two-tailed t-test). Moreover, 39% fewer AZs were seen to engage in both transmission modes, while a larger fraction of AZs solely responded to APs (**Figure 2 – figure supplement 3B**), indicating that AP-stimulation reduces spontaneous transmission at AP-evokable AZs. As the only difference between the two analyses is whether spontaneous activity is measured in isolation or in-between APs, our results clearly demonstrate that AP-evoked transmission reduces spontaneous transmission at BRP-positive AZs. Because this is unlikely due to saturation of NT receptors (see below) this cross-depletion indicates the use of common presynaptic resources, including a shared pool of SVs used for both transmission modes.

We previously found that both transmission modes were positively correlated to the AZ-levels of BRP and Unc13A (Reddy-Alla et al., 2017) and here sought to investigate whether a common molecular dependence extends further, to voltage gated Na^+^ and/or Ca^2+^ ion channels which trigger AP-evoked transmission. Application of the voltage gated Na^+^ channel blocker tetrodotoxin (TTX) did not affect the frequency of spontaneous events (**Figure 3A**), indicating their independence from these channels. Instead, GCaMP imaging detected a strong reduction of spontaneous activity upon application of the voltage gated Ca^2+^ channel blocker CdCl_2_ (**Figure 3B**) (Ryglewski et al., 2012), which could implicate a dependence on these channels. However, the GCaMP signals themselves also tended to be affected (a lower average amplitude and slightly different kinetics of the GCaMP signals were seen, **Figure 3B**) which raises the concern that Cd^2+^ may interfere with optical signal detection. We therefore complemented our analysis with electrophysiological recordings, which -unlike our optical analysis at BRP positive AZs of 1b boutons-sample spontaneous and AP-evoked events across the entire NMJ.

**Figure 3.**
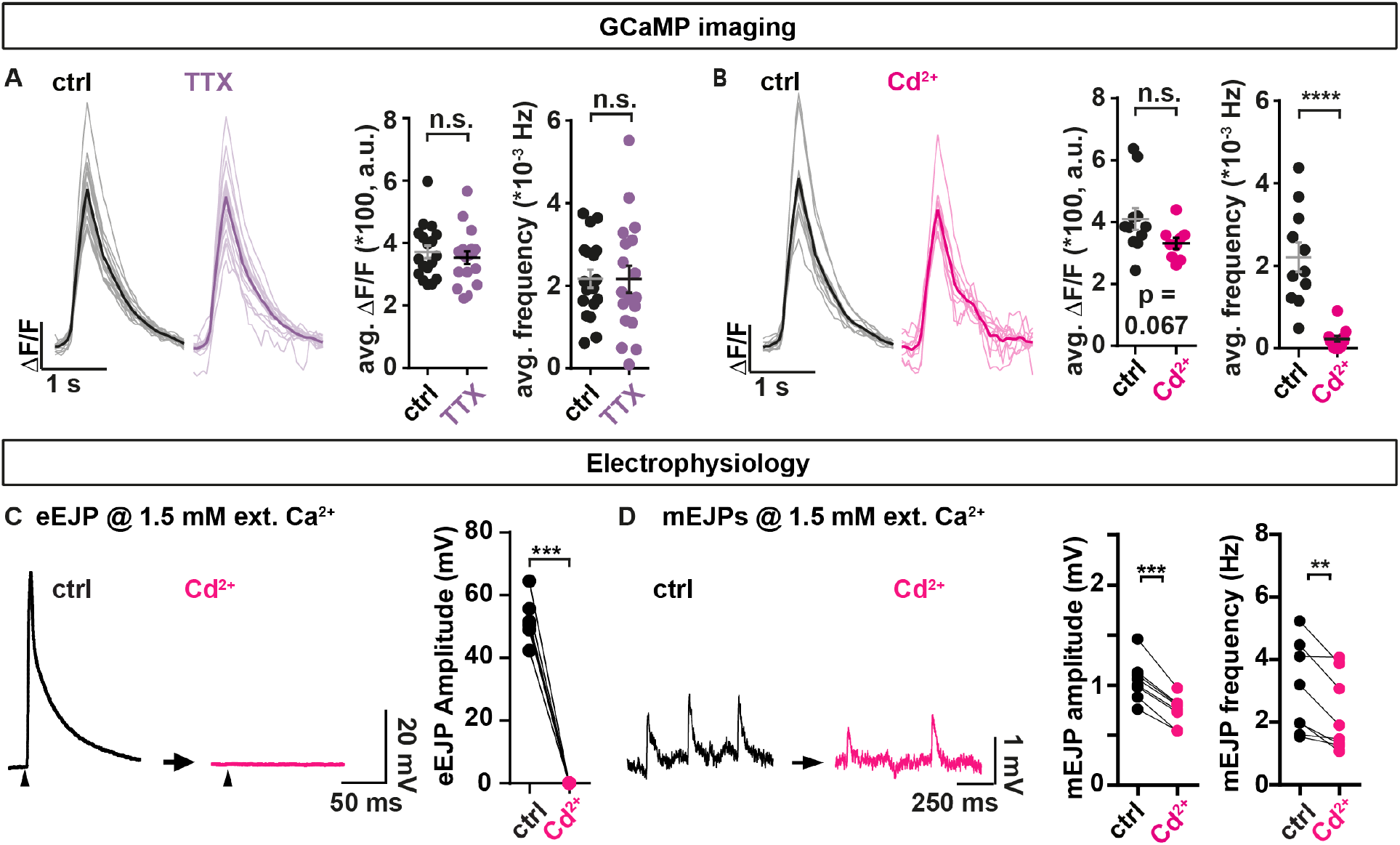
The non-specific voltage gated Ca^2+^ channel blocker Cd^2+^ affects spontaneous transmission. (**A&B**) analysis of spontaneous transmission using GCaMP-imaging at BRP-positive AZs to study the consequences of the voltage gated Na^+^ channel blocker TTX (purple, 1 µM) and the Ca^2+^ channel blocker Cd^2+^ (magenta, 740.7 µM). Left: Single cell mean (faint) and cell-wise average (solid) traces of spontaneous events. Right: Cell-wise quantification of GCaMP event amplitudes and event frequencies. (**C&D**) Assessment of the consequences of Cd^2+^-application (300 µM, black→magenta) on AP-evoked eEJPs (C) and on spontaneous mEJPs (D) in paired current clamp recordings of muscle 4 NMJs. (C) Left: Representative eEJP before (black) and after (magenta) Cd^2+^ application. Arrowheads indicate time of stimulation. Right: Animal-wise quantification of eEJP amplitudes before (black) and after (magenta) Cd^2+^ application. (**D**) Left: Representative mEJC traces before (black) and after (magenta) Cd^2+^ (300 µM) application. Right: Cell-wise quantification of mEJC amplitudes and -frequencies. Number of animals in (A): N(ctrl) = 18; N(TTX) = 18. Number of animals in (B): N(ctrl) = 11; N(Cd^2+^) = 11. Number of animals in (C&D): N=8. Horizontal lines in (A&B) indicate mean, error bars SEM. n.s. not significant; * p<0.05; *** p<0.001; **** p<0.0001. Two-tailed parametric Student’s t-test for comparisons in A&B. Paired parametric t-test for comparisons in (C&D). See also **Figure 3 – figure supplement 1&-2**.

Current clamp recordings without current injection into the muscles monitor synaptic activity while allowing the muscle’s membrane potential to vary naturally with the transmission (similar as in our optical recordings). Paired analyses of AP-evoked transmission at muscle 4 NMJs revealed that Cd^2+^ fully blocked AP-<u>e</u>voked <u>E</u>xcitatory <u>J</u>unction <u>P</u>otentials (eEJPs) (**Figure 3C**). Comparison of spontaneously occurring <u>m</u>iniature <u>E</u>xcitatory <u>J</u>unctional <u>P</u>otentials elicited by single SV fusion events before and after treatment revealed a reduction in both their amplitude and frequency by Cd^2+^ (**Figure 3D**). This effect appeared specific to a blockage of Ca^2+^ influx by Cd^2+^, as Cd^2+^ had no effect on these measures in animals recorded in the absence of extracellular Ca^2+^ **Figure 3 – figure supplement 1A** (and we excluded run-down as responsible for decreased mEJP frequencies in this paradigm using a mock treatment) (**Figure 3 – figure supplement 1B**). Thus, Cd^2+^ treatment clearly alters spontaneous transmission. However, whether the reduced mEJPs frequency was (entirely) due to a decrease in spontaneous presynaptic NT release could not be discerned with certainty due to the simultaneous decrease in mEJP amplitudes (which could have reduced some events below our detection limit).

Decreased mEJP amplitudes could be due to a Cd^2+^-dependent depolarization of the muscle’s resting membrane potential. We therefore additionally performed voltage clamp recordings where this potential is set by the experimenter. In these we monitored spontaneous and AP-evoked release by measuring the currents required to clamp this potential. This revealed a decrease in the number of spontaneous events without a change of their amplitudes (**Figure 3 – figure supplement 2**), thereby alleviating the concern that the decrease in spontaneous event frequency is secondary to a in their amplitudes. Together, our electrophysiological analysis confirms a partial dependence of spontaneous NT release on voltage gated Ca^2+^ channels. However, the observed reduction of events following Cd^2+^ application was much smaller than in the optical assay (compare **Figure 3B** and **Figure 3 – figure supplement 2**) which we attribute to a large proportion of spontaneous NT release events from non-BRP positive locations (see discussion).

While above experiments point to the use of common presynaptic resources, another question pertains to the postsynaptic receptors that detect neurotransmitters that could either be segregated or shared between transmission modes (**Figure 4A,B**). Due to the strong predictive value of spontaneous activity for AP-evoked responses, spontaneous transmission might serve as a continuous neural signal to monitor AP-sensitivity. If so, this “mixed” channel should activate the same postsynaptic NT receptors (**Figure 4B**). To test this, we stimulated AP-evoked NT release in the presence of the use-dependent glutamate receptor blocker Philanthotoxin (PhTx) (Frank et al., 2006; Sara et al., 2011) and quantified whether this affected spontaneous transmission (**Figure 4C**), which only happens if receptors are shared (**Figure 4A,B**).

**Figure 4.**
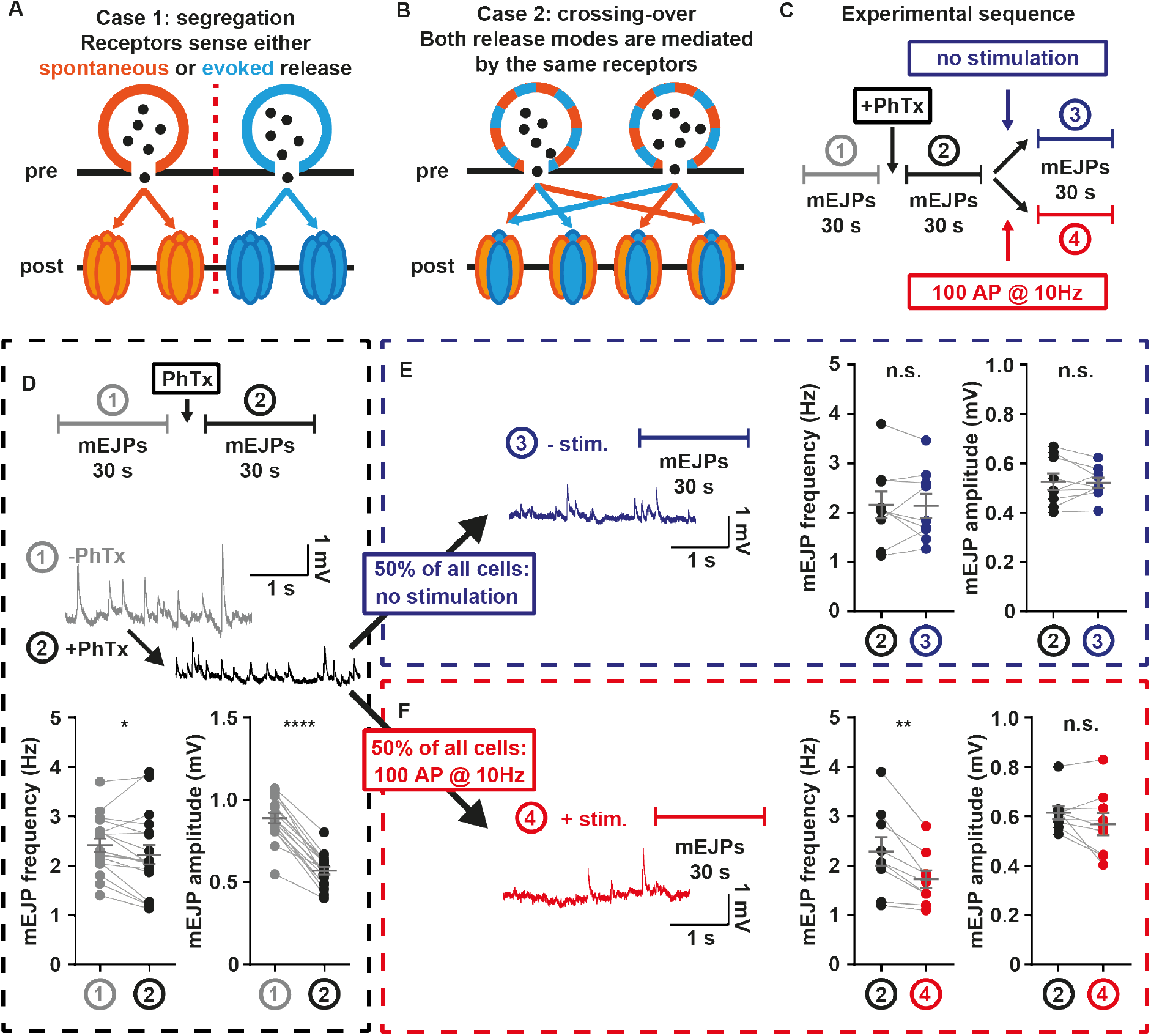
Postsynaptic receptors share sensitivity for both release modes. (**A**) Illustration case 1: Spontaneous (orange) and AP-evoked (blue) transmission rely on distinct postsynaptic receptors. (**B**) Illustration case 2: Both release modes use the same postsynaptic receptors. (**C**) Experimental setup: After 30 s mEJP baseline recordings, larvae are treated with 4 µM PhTx, and spontaneous mEJPs are recorded for another 30 s. Then, one half of animals (measurement 4, red) undergoes stimulation at 10 Hz over 10 s, the other half (measurement 3, blue) receives no stimulation. Another 30 s of mEJPs are recorded, yielding traces as shown in (D-F). (**D**) mEJP frequency and amplitude quantification before (grey; measurement 1) and after (black; measurement 2) PhTx application (N = 18 animals) (**E**) mEJP frequency and amplitude quantification after PhTx application and without stimulation (N = 9 animals) (**F**) mEJP frequency and amplitude quantification after PhTx application and with stimulation (N = 9 animals). Means are shown, error bars indicate SEM. n.s. not significant; * p<0.05; ** p<0.01; **** p<0.0001, paired t-test See also **Figure 4 – figure supplement 1**.

We initially monitored baseline spontaneous transmission in electrophysiological current-clamp experiments for 30 s before and after PhTx application, which expectedly reduced the amplitude of postsynaptic “miniature” excitatory junction potentials (mEJPs) caused by spontaneous NT release (**Figure 4D**)(Frank et al., 2006). In half of the animals, the efferent nerve was then stimulated with 100 APs (10 Hz, **Figure 4F**), a stimulation suitable to block glutamate receptors in the presence of PhTx (**Figure 4 – figure supplement 1**). The other half of the animals received no AP stimulation (but saw a corresponding 10 s wait) (**Figure 4E**). In both groups, mEJPs were recorded for another 30 s. If spontaneous and AP-evoked activity were exclusively sensed by distinct postsynaptic receptors (**Figure 4A**), AP-stimulation should not affect spontaneous transmission. Contrasting this, a clear decrease of mEJP frequency selectively occurred in the group receiving the AP stimulations (**Figure 4F**). The effect was specific to the use-dependent block by PhTx, as stimulation alone had no effect (**Figure 4 – figure supplement 1A-C**), arguing against SV pool depletion or receptor desensitization as underlying cause. Our results show that receptor block induced by AP-evoked activity affects spontaneous neurotransmission, clearly indicating that both transmission modes activate the same receptors at AP-evocable synapses. This does not exclude the existence of additional “distinct” communication channels (i.e. ones dedicated to spontaneous transmission) but demonstrates that both transmission modes share NT receptors at AP-responsive AZs.

## Discussion

Our data demonstrate that the spontaneous activity of individual AP-evocable AZs is a predictor of their responsiveness to AP stimuli. This functional property could enable synapses to continuously monitor the evocable signal strength of a connection at rest, using highly uniform, spontaneous signals (“pings”) and would allow for detection and homeostatic compensation of dysbalances even before the connection is stimulated. Indeed, presynaptic homeostatic potentiation at the *Drosophila* NMJ increases NT release to compensate reduced postsynaptic NT receptor sensitivity by entirely relying on spontaneous (not AP-induced) activity (Frank et al., 2006). Such mechanisms may even be more important for sparsely activated synaptic connections of the central nervous system and indeed this homeostasis also exists in the mammalian brain (Delvendahl et al., 2019).

The easiest explanation for the high interdependence of both activity modes at AP-responsive AZs is a reliance on the same machinery. We find that both transmission modes activate shared NT receptors because AP-stimulation in the presence of the use-dependent PhTx (but not AP-stimulation alone) decreased spontaneous transmission (**Figure 4, Figure 4 – figure supplement 1A-C**). A previous study using PhTx at the *Drosophila* NMJ concluded no effect of AP-dependent NT receptor block on spontaneous transmission (Peled et al., 2014). However, those experiments were compared across different groups of animals (which is less sensitive) and were performed over far longer times (25 min vs. 10 s here) during which compensatory, homeostatic mechanisms take place with this treatment (Frank et al., 2006; Davis and Müller, 2015; Harris and Littleton, 2015).

We show that AP-stimulation reduces spontaneous activity (**Figure 2 – figure supplement 3B**). This could indicate the saturation or depletion of a common pre- or postsynaptic resource. It seems unlikely that this effect is caused by the saturation of postsynaptic NT-sensing receptors because even stronger AP-stimulation (100 APs @ 10 Hz) elicited no detectable effect on the rate of spontaneous transmission in electrophysiological recordings (**Figure 4 – figure supplement 1A-C**). Instead, it is more likely that the AP-induced reduction of spontaneous activity is due to a depletion of a common pool of SVs for both transmission modes (Groemer and Klingauf, 2007; Hua et al., 2010; Wilhelm et al., 2010). This implies a long-lasting inhibition of spontaneous activity recovering much slower rate of pool refilling than current estimates (Miki et al., 2016; Kobbersmed et al., 2020). The precise reason for this is unknown but may reflect a difference in such experiments that typically engage repetitive stimuli vs. the sparse stimulation used here (0.2 Hz).

Previous analyses indicated that both spontaneous and AP-evoked single-AZ activity were positively related to the local levels of BRP and Unc13A (Reddy-Alla et al., 2017). The common requirement on presynaptic proteins likely extends further as mutations of specific residues of SNAP25 or Syaptotagmin-1 (which both mediate AP-evoked activity) reduced both transmission modes (Xu et al., 2009; Weber et al., 2010). We here investigated whether a common dependence at the *Drosophila* NMJ also extended to voltage gated ion channels. While blocking voltage gated Na^+^ channels with TTX (which abolishes AP-evoked release) did not affect spontaneous transmission, we saw a strong reduction spontaneous transmission upon application of the non-specific voltage gated Ca^2+^ channel blocker Cd^2+^ in our GCaMP experiments (**Figure 3B**). This could mean that stochastic gating of VGCCs triggers spontaneous transmission by activating the same fusion machinery at AP-evocable AZs. Indeed, such a reliance of spontaneous activity on VGCCs has been demonstrated for other systems (Shahrezaei et al., 2006; Goswami et al., 2012; Ermolyuk et al., 2013). However, whether this is also the case for *Drosophila* synapses is debated. An argument against this is that unlike in other systems electrophysiologically measured spontaneous release rates at the *Drosophila* NMJ do not depend on the external Ca^2+^ concentration (Huntwork and Littleton, 2007; Xu et al., 2009; Groffen et al., 2010; Goel et al., 2017). However, one reason for this discrepance could relate to the locations of the spontaneous events predominantly detected using either approach.

In our GCaMP imaging analysis we specifically focus on spontaneous transmission from BRP-positve AZs, which are predominantly AP-resposnive (**Figure 2 – figure supplement 1**), therefore represening a “mixed channel” with both transmission modes. In contrast, electrophysiological recordings assess spontaneous transmission across the entire NMJ and therefore can additionally detect spontaneous transmission from synaptic contacts with no (or non-detectable) BRP. Indeed, our electrophysiological recordings demonstrated a much smaller inhibition of spontaneous activity by Cd^2+^ than seen with GCaMP imaging (**Figure 3B&D, Figure 3 – figure supplement 2**). This much reduced sensitivity could imply that synapltic connections without (or with low) BRP form a “dedicated spontaneous” communciation channel which might dominate overall spontaneous transmission at the NMJ. In fact, these (“non-BRP localized”) events could well represent the “dedicated communication channel” for spontaneous transmission shown to utilize distinct SVs, NT receptors, SNAREs (vti1a, VAMP7) and Ca^2+^ sensors (doc2b) and therefore might not be prone to manipulation of the AP-evoked machinery (Sara et al., 2005; Fredj and Burrone, 2009; Groffen et al., 2010; Ramirez et al., 2012; Bal et al., 2013). This may also explain some of the differences (opposite dependence on BRP, inverted correlation of signals) we observe to the study by Peled and colleagues who had studied GCaMP events throughout the NMJ in a null mutant of Rab3 where the spatial segregation of BRP-positive and BRP-negative areas is augmented (Graf et al., 2009; Peled et al., 2014).

What may determine the ratio of spontaneous events in the “mixed” or “separate” communication channel? This could, for instance, relate to a developmental trajectory, with nascent synapses forming the “dedicated spontaneous channel” where a specialized machinery first mediates the spontaneous NT release needed for maturation which at later times leads to the accumulation of the AP-responsive release machinery, generating the “mixed channel” (Truckenbrodt and Rizzoli, 2014; Walter et al., 2014; Akbergenova et al., 2018). In that case, the ratio between the two channels might greatly depend on the maturity of the model system and indeed spontaneous activity decreases and evoked transmission increases during maturation of neural cultures (Andreae et al., 2012). Thus, some of the differences reported in the literature regarding the in- or interdependence of transmission modes may rely on different states of maturation of the systems in which experiments were performed (e.g. between cultured neurons, brain slices or NMJs). Likewise, genetic deletion of synaptic components of the AP-evoked machinery may impede this transition in addition to their effect on communication along the “mixed” channel thereby increasing the proportion of events in the “dedicated spontaneous” channel. This may partly explain the divergent effects on both transmission modes upon null-mutation of genes encoding the synaptic proteins Synaptobrevin, Synaptotagmin-1, complexin or voltage gated Ca^2+^ channels (Littleton et al., 1993; Broadie et al., 1994; Geppert et al., 1994; Broadie et al., 1995; Deitcher et al., 1998; Reim et al., 2001; Schoch et al., 2001; Huntwork and Littleton, 2007; Hou et al., 2008; Martin et al., 2011; Yang et al., 2013). Thus, while our study identifies shared resources and a clear predictive role of spontaneous transmission for AP-evoked transmission in the “mixed channel”, the functional relevance of the relation between “dedicated spontaneous” and “mixed” transmission channels for synapse development, maintenance, and information transfer demand further investigation.

## Materials & Methods

### Resources Table

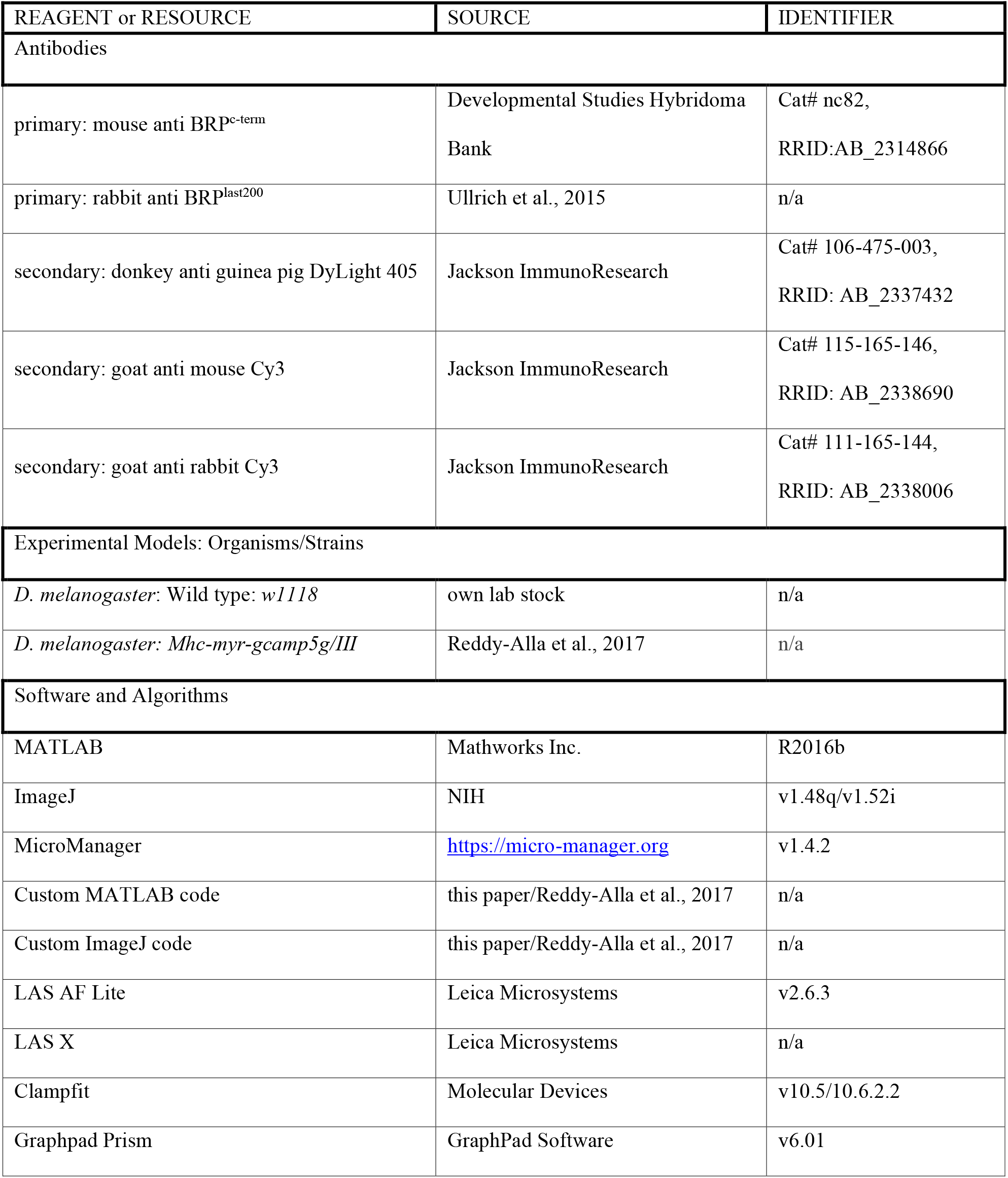
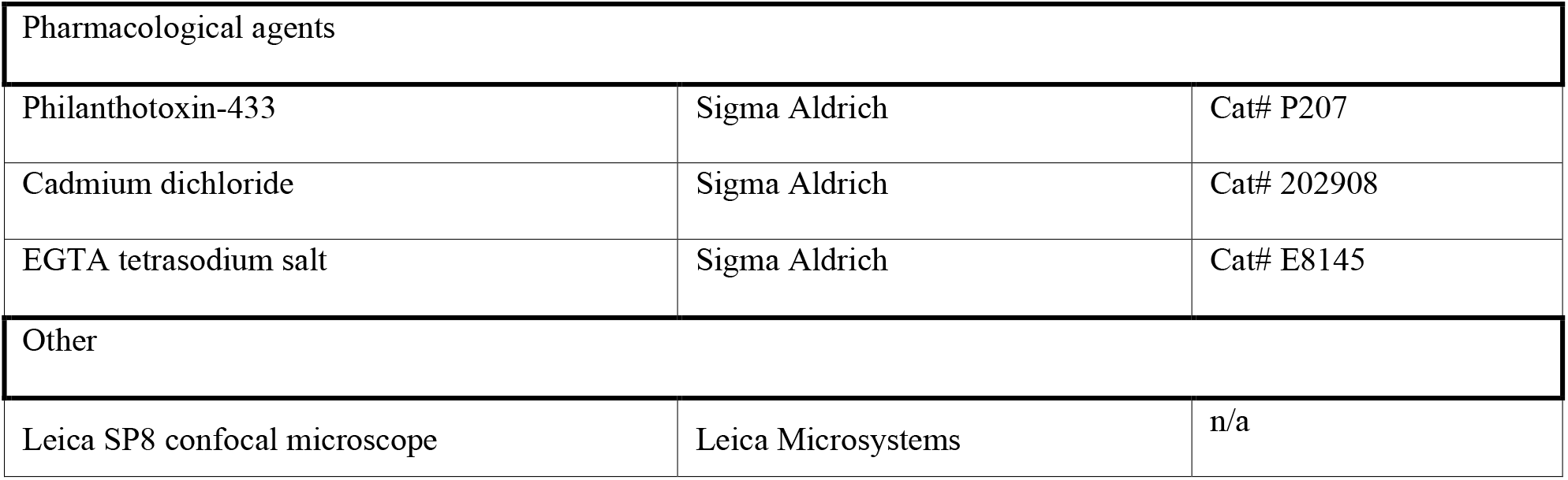

### Animal rearing and preparation

Experiments making use of genetically modified invertebrate animals have been registered with and approved by the respective authorities (Landesamt für Gesundheit und Soziales LaGeSo, Berlin), and were performed in accordance with German laws and regulations on biosafety. Animals were bred and maintained at standard laboratory conditions (Sigrist et al., 2003) on semi-defined fly medium (Bloomington recipe). Male and female flies were used for all experiments. Wild type *w1118* flies were used for the electrophysiology experiments shown in **Figure 3C&D, Figure 3 – figure supplement 1&2, Figure 4, and Figure 4 – figure supplement 1**. The following fly strain was used for all other experiments: *Mhc-myrGCaMP5G/+;* (Reddy-Alla et al., 2017). Third instar larvae were dissected as described in Qin et al., 2005 in standard Ca^2+^-free, hemolymph-like solution HL-3 (composition in mM: 70 NaCl, 5 KCl, 20 MgCl_2_, 10 NaHCO_3_, 5 Trehalose, 115 Sucrose, 5 HEPES; Stewart et al., 1994; low Mg^2+^-solution used for PhTx-electrophysiology experiments contained 10 mM MgCl_2_), adjusted to pH=7.2. Dissection was performed using fine insect pins on Sylgard-184 (Dow Corning, Midland MI, USA), by opening the dorsal body wall from posterior to anterior, removing all internal organs, and severing the motoneurons from the CNS without damaging the underlying body wall muscles, then removing the brain. For experimentation, the dissected larvae were then transferred to recording chambers on the respective recording setups, as detailed in the sections explaining electrophysiology and live calcium imaging.

### Electrophysiology

All electrophysiological experiments were performed at room temperature using sharp glass electrodes (borosilicate glass with filament, 0.86×1.5×80 nm, Science products, Hofheim, Germany) pulled with a Flaming Brown Model P-97 pipette puller (Sutter Instruments, CA, USA). Stimulating suction electrodes were pulled on a DMZ-Universal Puller (Zeitz-Instruments GmbH, Germany) and fire polished using a CPM-2 microforge (ALA Scientific, NY, USA). Recordings were performed in current clamp mode at muscle 6 (PhTx experiments) / muscle 4 (Cd^2+^/TTX experiments) in segments A2/A3 as previously described (Zhang and Stewart, 2010) using an Axon Digidata 1550A digitizer, Axoclamp 900A amplifier with HS-9A x0.1 headstage (Molecular Devices, CA, USA) and on a BX51WI Olympus microscope with a 40X LUMPlanFL/IR water immersion objective. Sharp intracellular recording electrodes with a resistance of 20-35 MΩ were made and back-filled with 3 M KCl. Only cells with membrane potentials below -60 mV (PhTx experiments) / -40 mV (Cd^2+^ experiments) and membrane resistances greater than 4 MΩ were considered. Recordings were acquired using Clampex software (v10.5) and sampled at 10-50 kHz, filtering with a 5 kHz low-pass filter. Analysis for all electrophysiological datasets was performed with Clampfit (v10.5/10.6.2.2) and Graphpad Prism 6 software. mEJPs were further filtered with a 500 Hz Gaussian low-pass filter. A single mEJP template was generated for each cell and used to identify individual mEJPs, and to calculate the mean mEJP amplitude and frequency per cell.

### Current clamp experiments to determine VGCC role in spontaneous SV release

Current clamp recordings using CdCl_2_ in order to block VGCCs were performed at room temperature from muscle 4 of abdominal segments A2-A4 (**Figure 3C&D, Figure 3 – figure supplement 1**) using 2 ml standard HL3 medium containing either 1.5 mM CaCl_2_ **(Figure 3C&D, Figure 3 – figure supplement 1C&D)** or 0 mM CaCl_2_ together with 2 mM of the Ca^2+^ chelator EGTA-tetrasodium salt (Sigma, Germany, E8145) to buffer residual Ca^2+^ traces **(Figure 3 – figure supplement 1A&B)**. Recordings shown in **Figure 3C&D, Figure 3 – figure supplement 1** were obtained before and after the addition 300 µM CdCl_2_ (magenta) or the equivalent volume of dH_2_O as control (grey) (see pharmacology section) in a strictly paired fashion. In detail, starting with a CdCl_2_-free extracellular medium (“ctrl”) a single AP was evoked in motoneurons (8 V, 300 µs pulse) using an ISO-STIM 01D stimulator (NPI Electronic, Germany) followed by a 30 s resting period. Sequentially, spontaneous mEJPs were recorded for 30 s followed by an immediate exchange of 1 ml bath solution (2 ml total bath volume) by 600 µM CdCl_2_-HL3 (“Cd^2+^”) or dH_2_O-HL3 (“ctrl”) within a resting period of 2 min. Afterwards, another single AP was evoked followed by 30 s rest and recording of 30 s spontaneous activity. eEJP amplitudes were determined by quantifying the maximal voltage deflection from the basline following an AP (values in the standard noise range were considered as zero).

### Two-electrode voltage clamp experiments to determine VGCC role in spontaneous SV release

TEVC recordings were performed at room temperature from muscle 4 of abdominal segments A2-A4 (**Figure 3 – figure supplement 2**). Signals were recorded using a 5 KHz low-pass filter at a sampling frequency of 20 KHz using the Digidata 1440A digitizer (Molecular devices, Sunnyvale, CA, USA) with Clampex (v10.6) software and Axoclamp 900A amplifier (Axon instruments, Union City, CA, USA) with Axoclamp software. Only cells with resting membrane potentials below -49 mV and membrane resistances above 4 MΩ prior to measurements were included in the datasets. TEVC recordings shown in **Figure 3 – figure supplement 2** were performed at 1.5 mM extracellular CaCl_2_. Recordings shown in **Figure 3 – figure supplement 2** were obtained in the presence of 740.7 µM CdCl_2_ (magenta) or the equivalent volume of dH_2_O as control (black) (see pharmacology section). Cells were clamped at a holding potential of -55 mV.

### Current clamp experiments to determine receptor sensitivity to different SV release modes

For current clamp experiments using PhTx to determine postsynaptic receptor field sensitivity to both release modes (**Figure 4**), the Sylgard block and completed larval preparation was placed in the recording chamber which was filled with 2 ml HL3 (0.4 mM CaCl_2_, 10 mM MgCl_2_). eEJPs were recorded by stimulating the appropriate nerve at 10 Hz, 100 times (8 V, 300 µs pulse) using an ISO-STIM 01D stimulator (NPI Electronic, Germany).

Spontaneous mEJPs for analysis shown in **Figure 4D-F** and **Figure 4 – figure supplement 1A-C** were recorded for 30 seconds. 1 ml of solution was then removed from the bath without disturbing the preparation or electrodes and 1 ml of HL3 added containing PhTx-433 (Sigma-Aldrich, MO, USA), mixing gently with the pipette to a final bath concentration of 4 µM PhTx. Spontaneous mEJPs were recorded immediately, again for 30 seconds. Stimulation was performed at 10 Hz for 10 seconds to measure eEJPs or, in the case of control recordings, 10 seconds passed without stimulation. Finally, mEJPs were recorded for 30 seconds. Recordings shown in **Figure 4 – figure supplement 1A-C** were performed as above, using HL3 lacking PhTx-433, as the exchange solution.

### Pharmacology

Philanthotoxin (PhTx-433) used for experiments in **Figure 4** was obtained from Sigma Aldrich (subsidiary of Merck KGaA, Darmstadt, Germany) and diluted to a stock concentration of 4 mM in dH_2_O. In experiments, it was used at a concentration of 4 µM in HL-3 by applying it directly to the bath (see electrophysiology method section). Tetrodotoxin-citrate (TTX) used for experiments shown in **Figure 3A** was obtained from Tocris (subsidiary of Bio-Techne, Minneapolis, MN, USA) and diluted in dH_2_O to a stock concentration of 1 mM. In GCaMP experiments, it was used at a concentration of 1 µM in HL-3 (5 µL 1 mM TTX/dH_2_O stock in 4.6 mL HL-3 and 0.4 mL dH_2_O (diluting the HL-3 components to (in mM): 64.4 NaCl, 4.6 KCl, 18.4 MgCl_2_, 9.2 NaHCO_3_, 4.6 Trehalose, 105.8 Sucrose, 4.6 HEPES)), and imaging began after 2 minutes of incubation time. Cadmium dichloride (CdCl_2_) was obtained from Sigma Aldrich. For the experiments shown in **Figure 3B** a stock solution with a concentration of 10 mM was made and for experiments further diluted to a final concentration of 740.7 µM in HL-3 (1.12 mL 10 mM CdCl_2_/dH_2_O stock in 14 mL HL-3 and a corresponding amount of 1.12 mL dH_2_O in 14 mL HL-3 controls, diluting the HL-3 components to (in mM): 64.8 NaCl, 4.63 KCl, 18.52 MgCl_2_, 9.26 NaHCO_3_, 4.63 Trehalose, 106.48 Sucrose, 4.63 HEPES); measurements began after 2 minutes of incubation time. The Current Clamp experiments depicted in **Figure 3C&D** and **Figure 3 – figure supplement 1** were performed at a later time. For these, a different stock solution was made with a concentration of 300 mM in dH_2_O, and used at a final concentration of 300 µM in HL-3 for electrophysiological experiments. Control experiments shown in **Figure 3 – figure supplement 1C&D** were performed using the same amounts of the respective solvent (CdCl_2_: dH_2_O/HL3, PhTx: dH_2_O).

### Live Calcium-Imaging

GCaMP live imaging experiments were conducted in 2 mL HL-3 containing 1.5 mM CaCl_2_ (except for Ca^2+^-titration in **Figure 1 – figure supplement 1H**: 0.4, 0.75, 1.5, 3, 6, 12 mM) on an Olympus BX51WI epifluorescence microscope, using a water immersion LUMFL 60x 1.10 w objective. A Lambda DG-4 (Sutter Instrument Company, Novato CA, USA) white light source was used to illuminate the samples through a GFP excitation/emission filter set. For experiments in **Figure 1 – figure supplement 1H** and **Figure 2 – figure supplement 1D-H**, a newer light source of the same model was used in combination with an Olympus ND25 neutral density filter. Images were acquired in camera-native 16-bit grayscale using an Orca Flash v4 sCMOS camera (Hamamatsu Photonics, Hamamatsu, Japan) under constant illumination with an exposure of 50 ms per frame, resulting in an effective imaging frame rate of 20 Hz. For all GCaMP analysis, spontaneous events were recorded from 1b NMJs in muscle 4, abdominal segments 2-4, for 100 s (120 s in the case of TTX and Cd^2+^ experiments shown in **Figure 3A&B**). For experiments involving the imaging of AP-induced (‘evoked’) events, the efferent motoneuronal axon bundle innervating the same muscle was sucked into a polished glass capillary containing bath HL-3. The glass capillary was held in place by a patch electrode holder (npi electronic, Tamm, Germany), and contained a chlorided silver wire electrode, which connected to a pipette holder (PPH-1P-BNC, npi electronic, Tamm, Germany). After recording of spontaneous events, 36 single stimuli were applied as a square depolarization pulse of 300 µs at 7 V, 0.2 Hz for 180 s using a connected S48 stimulator (Grass Technologies, now part of Natus Medical Inc., Pleasanton, CA, USA), except for analysis shown in **Figure 2 – figure supplement 2**, where the experimental sequence was reversed. Imaging start/end was controlled by µManager software (version 1.4.20, https://micro-manager.org), and stimulation was administered through software (Clampex 10.5.0.9, Molecular Devices, San Jose, CA, USA) controlling a DA/AD converter (DigiData 1440A, Molecular Devices, San Jose, CA, USA). All videos acquired in 16-bit were then converted to 8-bit using ImageJ (version 1.48q). See section Image processing and analysis for further procedures and details.

### Immunohistochemistry

After live imaging experiments, larval tissue was fixated for 10 min at RT using fresh 4% PFA in 0.1 mM PBS. Fixated samples (max. 8 per 1.5 mL sample cup) were then stored in 1 mL 1xPBS until all samples had been collected, but 6 hours at most. Then, off-target epitope blocking was performed in 1xPBS containing 0.05% Triton-X100 (PBS-T) and 5% normal goat serum (NGS) (total volume: 1000 µL) for 45 min on a wheel at RT, 17 rpm. Immediately after, the mix was replaced by an identical mixture and the respective first antibody was added at the following concentrations: mouse BRP^C-term^ (1:1000, Developmental Studies Hybridoma Bank, University of Iowa, Iowa City, IA, USA), rabbit BRP^last200^ (1:1000) (Ullrich et al., 2015). Samples were incubated with the primary antibody overnight (15-16 h) at 4°C on a sample wheel. Afterwards, samples were washed four times in PBS-T for 30 min at RT. Secondary antibodies were applied (4 h, RT) in PBS-T containing 5% NGS at the following concentrations: donkey anti guinea pig DyLight 405 (1:500, Jackson ImmunoResearch, West Grove, PA, USA), goat anti mouse Cy3 (1:500, Jackson ImmunoResearch), goat anti rabbit Cy3 (1:500, Jackson ImmunoResearch, West Grove, PA, USA). After this, they were washed with PBS for 30 min and finally mounted on 26×76 mm glass slides (Paul Marienfeld GmbH, Lauda-Königshofen, Germany) in VectaShield (Vector Laboratories, subsidiary of Maravai Life Sciences, San Diego, CA, USA) under 18×18 mm cover glass slides (Carl Roth GmbH, Karlsruhe, Germany) using clear nail polish to seal off the sides of the cover glass slide. The samples were then stored at 4°C and imaged within a week as described in the confocal microscopy and image processing section.

### Confocal microscopy and image processing

Confocal imaging of immunohistochemically stained samples was performed on a Leica SP8 confocal quadruple solid-state laser scanning system (excitation wavelengths: 405, 488, 552, 635 nm), and operating on LAS X software (Leica Microsystems, Wetzlar, Germany) with a 63x 1.4 NA oil immersion objective at room temperature. Pixel edge length was 100 nm at a zoom factor of 1.8 and a z-step size of 0.5 µm for all datasets. Care was taken to choose fluorophores with non-overlapping excitation/emission spectra (see immunohistochemistry section), and confocal GCaMP images were always acquired without additional IHC at 488 nm excitation. Single z-stack images from all channels were exported from the proprietary .lif-format into TIF images using LAS AF Lite software (version 2.6.3, Leica Microsystems, Wetzlar, Germany) and converted to 8-bit grayscale maximal projections using custom ImageJ/Fiji software (version 1.52i, available upon request).

### Image processing and analysis

#### Stabilization of live-imaging videos

As further analysis of GCaMP live-imaging videos was highly reliant on a stable position of the NMJ over time, all 2D-translational movement (in x,y-direction) of the muscle during the recording had to be corrected for. This was done as shown in Reddy-Alla et al., 2017, and is described in the following. Converting videos of mhc-myr-GCaMP5G from 16-bit to 8-bit grayscale was done in ImageJ.

After conversion from 16-bit to 8-bit, the 8-bit multipage .TIF-video file (‘stack’) was loaded into MATLAB and a subregion of the first frame, containing the whole GCaMP-positive 1b NMJ, was chosen as a reference for the registration process using the MATLAB function *getrect*. Using the MATLAB function *normxcorr2*, every subsequent frame was then 2D-translated by a simple *x,y*-shift until the highest cross-correlation between pixel values of the current frame and the first frame was achieved. This procedure was repeated for all pairs of the first frame and succeeding frames. For this procedure, all images were Gaussian filtered (MATLAB function *imgaussfilt*) with a sigma value of 5 for noise reduction.

### Alignment of confocal images to live-imaging videos

Next, we compensated for fixative-induced anisotropic deformation, orientation and size changes in confocal images by registering them to GCaMP-videos in ImageJ using the plugin “TurboReg” (Biomedical Imaging Group, EPFL, Switzerland; Thévenaz et al., 1998). An affine transformation that used three reference points in each image applied *x,y*-translation, rotation, and *x,y*-shearing where necessary to get the optimal overlay between GCaMP-signal in confocal and live-imaging. In rare instances, a bilinear transform using 4 reference points was necessary to achieve sufficient overlay between confocal image and GCaMP video. The necessary transformation found for the confocal GCaMP image was then applied identically to all other channels.

### Quantification of single AZ activity-levels

In order to quantify protein and activity levels on the level of single AZs, we first defined ROIs in the confocal BRP channel by applying the ImageJ function *find maxima* using threshold values between 10 and 20. Circular ROIs with a diameter of 650 nm were centrally overlaid at all *x,y*-positions found with this procedure. The integrated density (sum of intensities of all pixels whose sub-pixel center lay within the borders of the circle) in all ROIs was then saved for each confocal frame and live-imaging frame in .xls-format for later analysis in MATLAB. Additionally, a file with all corresponding *x,y*-coordinates of all ROIs was saved as a text file. Further, to correct for unspecific background fluorescence decay due to photobleaching, we shifted all ROIs to a region without GCaMP fluorescence and generated another .xls-file containing the fluorescence values in these ROIs over time. These values were later subtracted from the corresponding fluorescence signal in the original ROIs.

We then loaded these files into MATLAB for further analysis. First, we determined all inter-AZ distances (all distances between every possible pair of AZs) using the squared Euclidean distance as shown in equation (1).

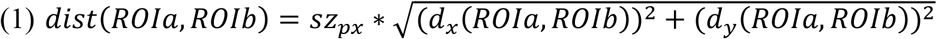

In equation (1), *ROIa* and *ROIb* are any of the determined ROIs, *sz*_*px*_ is the physical pixel edge length of 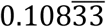 µm, and *d*_*x*_ and *d*_*y*_ are the vertical or horizontal pixel shift values in *x* or *y*, respectively, between both compared ROIs. This resulted in a diagonally symmetrical matrix of all possible inter-AZ distances. This distance was then used to exclude detecting another event within 2.5 µm (evoked activity measurements) or 1000 µm (spontaneous activity measurements) around one event in the same frame. We added another layer of security to exclude the detection of the same event twice by only considering the ROI with the highest amplitude within the given distance threshold and a single frame (each frame representing 20 ms of recording time).

The GCaMP fluorescence over time of each ROI was corrected for photobleaching as described before, by subtracting the fluorescence measured in the corresponding background ROI. We then performed a linear fit on each single fluorescence trace over time (separately for spontaneous and AP-evoked activity recordings), yielding two parameters reflecting its slope and y-intercept (MATLAB function *polyfit*). Using these parameters (slope *s* and y-intercept *int*), we performed a baseline correction for each time step *t* and each *ROI* as shown in equation (2).

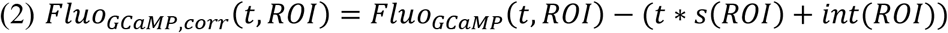

A custom procedure was then used to detect single peaks in the resulting fluorescence traces. All fluorescence traces were filtered by a 1D-filter using the MATLAB function *filter* (filter width: 5 frames). We then manually evaluated all instances in the fluorescence trace where the mean of the unfiltered signal over three consecutive frames exceeded a threshold of four times the SD of the filtered signal. As stated above, a circular distance threshold of 2.5 µm (AP-evoked activity measurements) or 1000 µm (spontaneous activity measurements) around each event was enforced to avoid unspecific detection of close by events in a single frame. When analyzing AP-evoked activity measurements, we only considered events that were detected within 1 s of the stimulus.

In order to generate activity maps as those shown in **Figure 2A**, we counted the number of detected events in each ROI and overlaid an inverted and contrast-adjusted IHC image of the respective protein at the NMJ with circles of corresponding sizes. The average signals shown in **Figure. 1F** were generated by averaging all detected events in each cell, and then averaging over all cell means.

### Spontaneous event detection between evoked events

Besides the “sequential” way of analyzing spontaneous activity measurements and then evoked activity measurements as described above, we also quantified spontaneous events that happened between stimuli (“interleaved”) as shown in **Figure 2 – figure supplement 3**. For this, we altered the procedure described above by one detail. While everything else happened as in our conventional approach to measure spontaneous activity, we suppressed the detection of evoked events and instead quantified SV release between stimuli by creating an exclusion list. This list included all time points 1000 ms after the stimulus, within which no fluorescence peaks would be considered as a signal.

### Survival analysis

A survival analysis quantifies the amount of “surviving” samples (in this case exclusively spontaneously active AZs) in face of an event that “kills” those samples (in this case trying to evoke release by a stimulus), i.e. switches them from one state to another over the course of the treatment. To analyze how many spontaneously active ROIs would “survive”, or maintain their exclusively spontaneous state by not showing any AP-evoked activity in the AP-evoked activity measurement (**Figure 2 – figure supplement 1**), we proceeded as follows. We loaded the results from the analysis of spontaneous and AP-evoked activity measurements (described above) containing all activity time points and AZ identities of spontaneous or evoked events into MATLAB. We then first found the number and identity of all AZs showing spontaneous activity. We created a data vector containing as many data points as there were frames in the AP-evoked activity recording (3600 over 180 s) and filled all positions with the number of AZs showing spontaneous activity we had found. Then, we found all time points of AP-evoked events in these AZs and subtracted 1 from the previously created vector at the time points of the evoked event to the end of the vector, resulting in a decreasing amount of exclusively spontaneously active AZs over the time of the AP-evoked activity measurement. For each cell, we then set the initial amount of exclusively spontaneously active AZs in that cell to 1 (100 %). Two models describing the mono-exponential survival decay were compared using Akaike’s information criterion (AIC; Akaike, 1974) to verify our approach as shown in equations (3) and (4), either excluding or including a plateau value of “surviving” exclusively spontaneously active AZs, respectively.

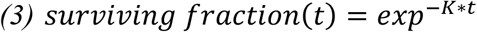

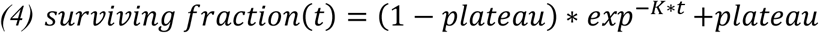

In equations (3) and (4), t is the timepoint at which survival is assessed, and K is the decay constant related to survival “half-life” (the timepoint at which half of the non-surviving AZs will have “died”) as 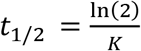. The plateau value represents the fraction of AZs that will not “turn” from spontaneous to mixed mode regardless of further stimulation. The difference in AIC warranted the use of the more complex equation (4).

### Analysis of relation between AP-evoked and spontaneous events

The response is number of evoked responses, which is analyzed with a generalized linear model using a negative binomial response distribution with a log link. This is a distribution for counts (a generalization of the Poisson distribution allowing for a flexible variance). The number of evoked events is modelled as a function of spontaneous events with a random effect of animal on the intercept (each animal has its own level). The number of spontaneous events enters as a factor.

To test for an increasing monotonic trend in the four estimated values of the mean of evoked events as a function of number of spontaneous events, we used the nonparametric Mann-Kendall trend test against a positive trend in a time series (Mann, 1945). The null is that there is no trend, so a small (one-sided) p-value suggests a positive monotonic trend. This is different from the test of an effect, which simply tests if the 4 estimated mean values can be assumed to be equal. The test was performed with the R-package trend, version 1.1.4, with option alternative = “greater” (Thorsten Pohlert, 2016). R package version 1.1.4. https://CRAN.R-project.org/package=trend

### Automated spontaneous event detection

For the automated detection of spontaneous vesicle fusion without respect to AZ positions shown in **Figure 1 – figure supplement 1B-F,H** we developed an additional set of custom MATLAB code. Single steps and results of the whole procedure on a single event are shown in **Figure 1 – figure supplement 2**. Stabilized 8-bit grey scale multipage .TIF-video files (**Figure 1 – figure supplement 2A**) were loaded into MATLAB, where the user could then manually select an area of the video with the NMJ of interest. Using the MATLAB function *bwboundaries*, a logical mask was then generated to find all pixels within the manually selected ROI. The chosen area was then extended by 20 pixels to each side, generating a rectangular selection taken from the original video. This cropped video was then further processed by slightly reducing noise using the *medfilt3* function (**Figure 1 – figure supplement 2B**), which smoothes noise in 3D arrays by taking the median grey value in a 3×3×3 pixel neighborhood. Next, the background was subtracted to leave only transient increases in fluorescence. For this, a maximum projection of 10 closely preceding frames was generated for every frame of the video, which was then subtracted from the current frame, where every resulting negative value was set to 0 (**Figure 1 – figure supplement 2C**). To avoid removing parts of an event, a ‘lag’ of 5 frames was included before the currently observed frame, resulting in a sequence of frames from the 15^th^ to 6^th^ before the current frame for the background subtraction. Every iteration of this process resulted in a single frame that was devoid of any basal GCaMP signal and excessive noise, only leaving transient fluorescence peaks that deviated from the brightest features of the last 15^th^ through 6^th^ frames. In addition to this background-subtracted video, another one was generated with the only difference being that here, instead of the maximum projection or 10 frames, an average projection of the same 10 frames was used to subtract the background. This video was then used for the exact determination of events by a 2D-Gaussian fit as described further down. A Gaussian filter (function *imgaussfilt* with a sigma of 3 pixels) was then applied to the resulting video for further noise removal (**Figure 1 – figure supplement 2D**). This was necessary for the next step, in which regions of connected (continuously bright) pixels above a threshold grey value of 2, and within the manual selection, were identified (**Figure 1 – figure supplement 2E**). For each of the identified regions, the median x,y-coordinates were found and temporarily defined as the location of the event (**Figure 1 – figure supplement 2F**). Detected events within 10 pixels of the edge of the video were removed, as they represented noise and were located outside the manual NMJ selection. A square 39×39 pixel region was then chosen around each event and a Gaussian fit was performed on a maximal projection of 6 frames (peak frame and 5 succeeding frames) of the second background-subtracted video, where the average of the 15^th^ to 6^th^ preceding frame was subtracted from each frame (**Figure 1 – figure supplement 2G**), as follows. A 2D-Gaussian was simulated (the ‘simulated image’, **Figure 1 – figure supplement 2H**) and fit to a maximal projection of six 39×39 px frames of an event (the ‘temporary image’) using equation (5):

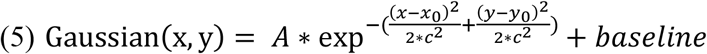

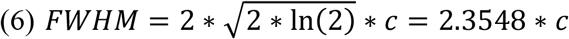

In equation (5), *x* and *y* are any of the coordinates on the image of the current event, *x*_*0*_ and *y*_*0*_ are the center coordinates of that image, *c* is a non-zero variable related to the full-width at half maximum (FWHM) of the Gaussian function as shown in equation (6), and *baseline* is a background correction factor. While the separation of x- and y-spread would allow non-symmetrical Gaussian fits, these parameters were kept identical in the fit, making the spread of the Gaussian uniform in 2D. An optimization procedure with the MATLAB function *fminsearch* was used to find the best parameters for the center x,y-coordinate of the Gaussian, its amplitude, its sigma value, and the baseline. An initial value of 20 was chosen for all five parameters. As a measure of the quality of the fit, a cost function was used that calculated the difference between the temporary image and the simulated image by subtracting them. As the success of *fminsearch* depends, among other factors, on the initial parameters, the optimization was additionally repeated three times with the best fit values of the previous run. The same optimization procedure with a genetic algorithm (which is less biased regarding initial parameters) yielded the same results at vastly longer processing times.

The analysis of spontaneous event amplitudes over increasing calcium concentrations shown in **Figure 1 – figure supplement 1H** was performed in *Mhc-myrGCaMP5G/+* larvae with the script described above at [Ca^2+^]_ext._ of 0.4, 0.75, 1.5, 3, 6 and 12 mM in HL3 by exchanging the external solutions between recordings in one animal and taking the cell-wise mean of GCaMP signals at their peak. The nonlinear fit on the cell-wise means was performed by assuming a Hill-relationship, where binding of Ca^2+^ to the sensor occurs with cooperativity *h*, and half-maximal fluorescence is reached at a concentration of [Ca^2+^]_ext._ of *K*_*A*_ as shown in equation (7). In that equation, *F*_*max*_ is the asymptotic maximal value of fluorescence at high [Ca^2+^]_ext._, and *C* is a baseline correction to allow a baseline fluorescence different from 0.

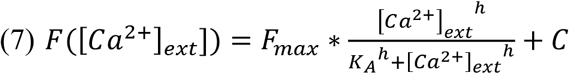

## Data and software availability

Source data is submitted for all figures. All ImageJ and MATLAB code used in this paper will be made available by the corresponding author (A.M.W.) upon reasonable request.

## Author contributions (CRediT taxonomy)

Conceptualization, A.T.G. and A.M.W.; Formal Analysis, A.T.G., A.W.M., M.J. and T.W.B.G.; Funding Acquisition, A.M.W.; Investigation, A.T.G., A.W.M., M.J., S.D., A.M.W. and T.W.B.G; Software, A.T.G. and A.M.W.; Supervision, A.M.W.; Visualization, A.T.G., A.W.M., M.J. and T.W.B.G; Writing – Original Draft, A.T.G. and A.M.W. Corresponding & Lead Author, A.M.W.

## Acknowledgments

We thank Stephan Sigrist and Volker Haucke for helpful discussion of the manuscript. We would like to thank Sabine Hahn for excellent technical assistance. M.J. was funded by a PhD-fellowship from the Einstein Center for Neurosciences Berlin. This work was funded by grants from the Deutsche Forschungsgemeinschaft (DFG, German Research Foundation): Project-ID 278001972, TRR 186 to A.M.W. and Project-ID 261020751, Emmy-Noether Programme to A.M.W.. This work was supported by the Novo Nordisk Foudation: Grant number: NNF19OC0056047, Young Investigator Award to A.M.W..

