## Supplementar Material for "Spontaneous neurotransmission at evocable synapses predicts their responsiveness to action potentials"

### Supplementary Figures

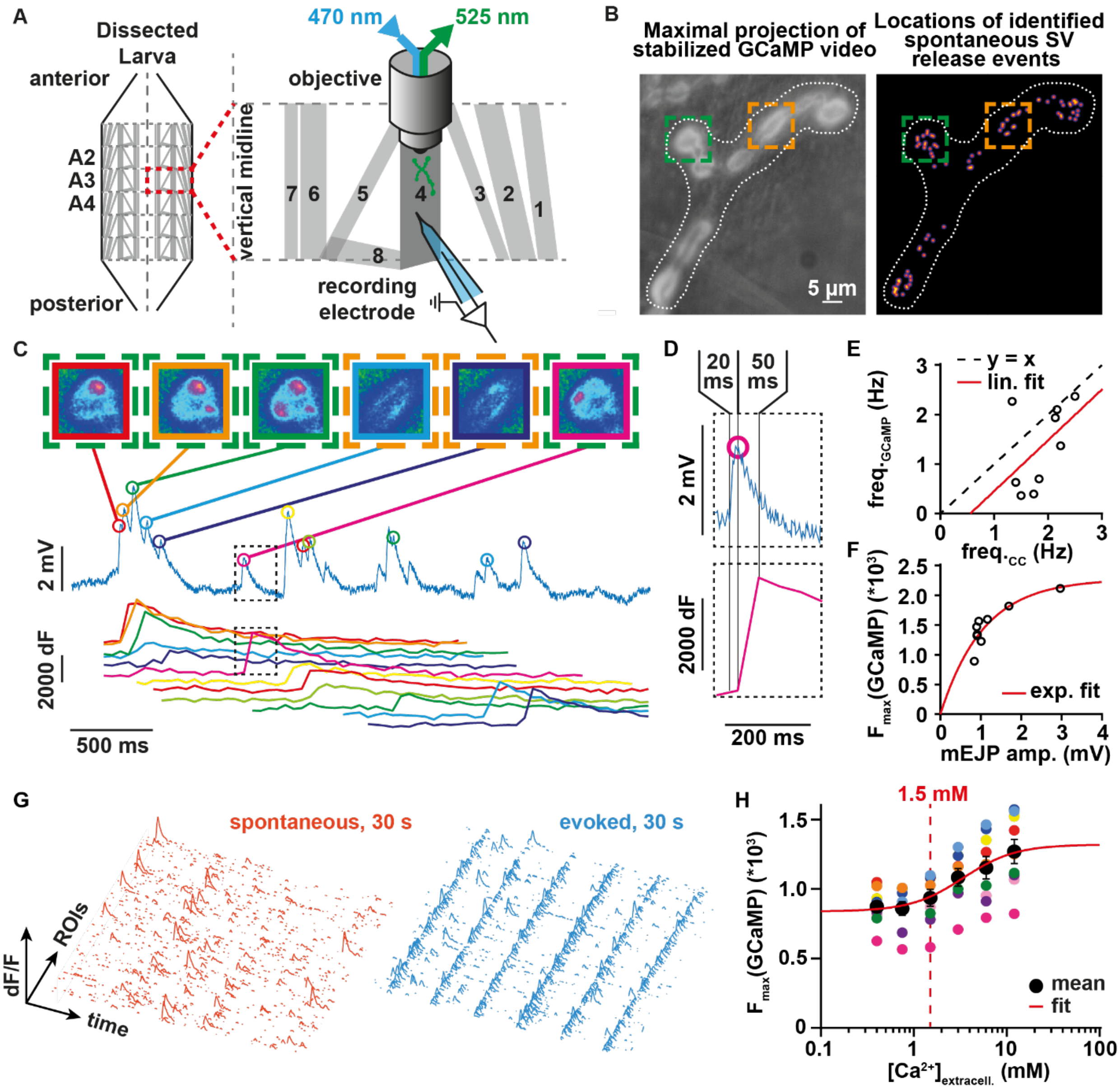

**Figure 1 – figure supplement 1. Further characterization of optical signals and their relation to synaptic activity.** (**A**) Scheme of the experimental setup; current clamp recordings and GCaMP fluorescence measurements are performed in the same muscle 4 NMJ. (**B**) Spontaneous event detection in GCaMP fluorescence assay independent throughout the NMJ (see Methods & **Figure 1 – figure supplement 2** for further details). Two ROIs over active boutons are marked in green and orange. (**C**) Events detected in current-clamp recordings and GCaMP fluorescence assay coincide to a large degree. Spontaneous events in bouton ROIs marked in **B** are observable with high spatial and temporal resolution (**D**) Blow-up of black ROI in **C** to show details of the temporal relation of current-clamp (top) and fluorescence (bottom) measurement. (**E**) Animal-wise (N = 9 cells&animals) spontaneous event frequencies measured in current-clamp recordings plotted against frequencies measured in fluorescence recordings. Linear fit on cell means in red, dashed black line represents x = y. (**F**) Animal-wise (N = 9 cells&animals) mEJP amplitudes measured in current-clamp plotted against maximal fluorescence amplitudes measured in fluorescence assay.

Exponential fit on cell means in red. (**G**) 3D representation of 30 s of spontaneous (orange) and AP-evoked (blue) event amplitudes over all ROIs in one NMJ (**H**) Quantification of spontaneous event amplitudes over six extracellular $Ca^{2+}$ concentrations (0.4, 0.75, 1.5, 3, 6, 12 mM) shows no saturation at physiological 1.5 mM $[Ca^{2+}]_{ext.}$ (N = 9 animals). Red line represents hill curve fit on individual values. Data is shown as mean amplitude per animal (colored dots) or animal-wise mean±SEM (black). Scale bar in **B**: 5 µm

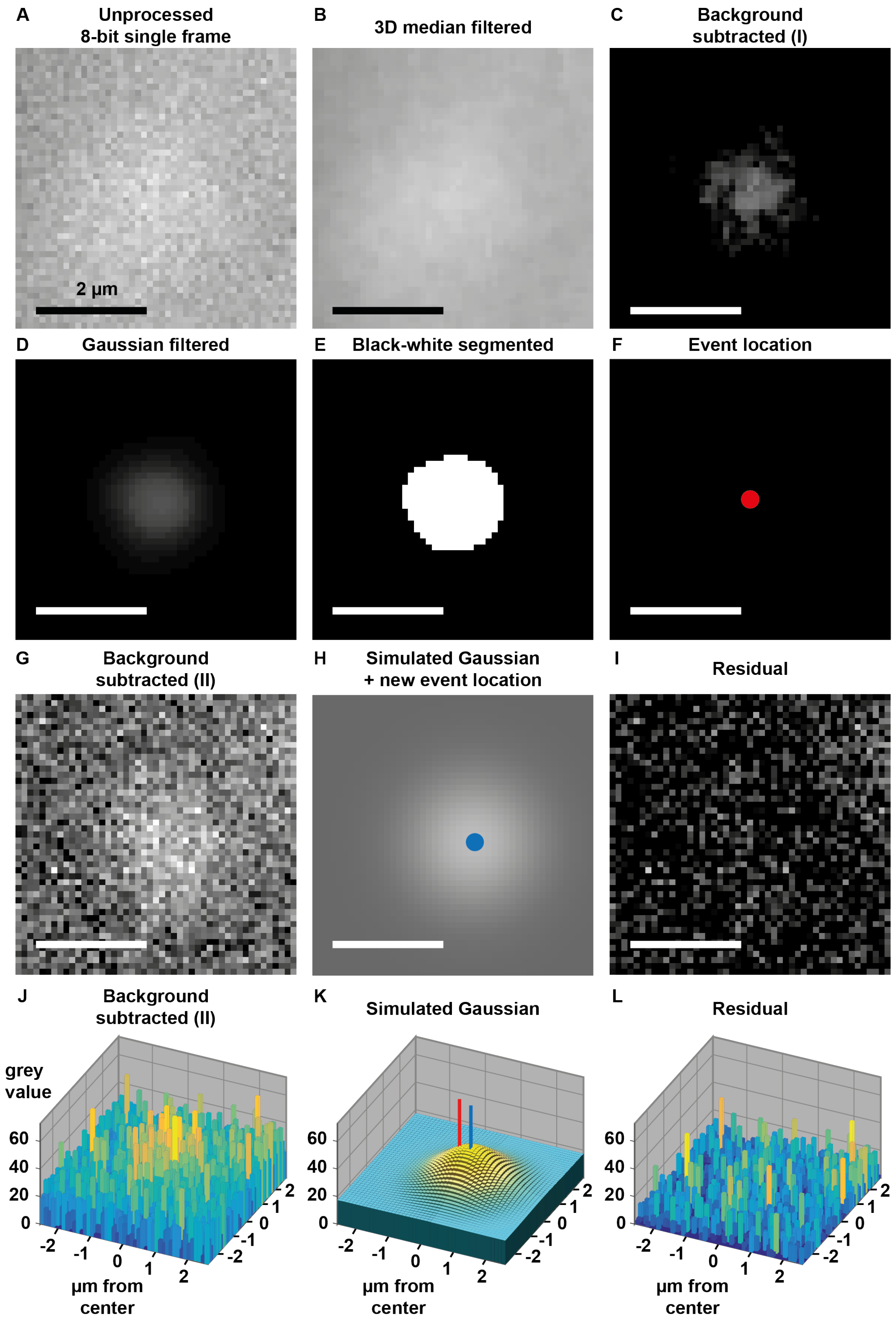

**Figure 1 – figure supplement 2. Sequence of event detection algorithm for spontaneous activity throughout the NMJ. (A)** 47x47 pixel cutout from the original, 8-bit video showing a typical spontaneous event. **(B)** Image from A after 3D median filtering for noise reduction. **(C)** Image from **B** after subtraction of brightest features of the 10$^{th}$

through $6^{th}$ preceding frames. **(D)** Image from C after application of a Gaussian filter for noise reduction. **(E)** Image from **D** segmented into grey values below or equal to 2 (black) or greater (white). **(F)** Determined location of the event. **(G)** Maximum projection of 6 frames from original video **A** after subtraction of average features of the $10^{th}$ through $6^{th}$ preceding frames. **(H)** 2D Gaussian fit to **G**. **(I)** Residual of Gaussian fit: Result of subtracting **H** from **G**. (**J)-(L)** 3D representations of images in **G-I**. Scale bar: 2 µm

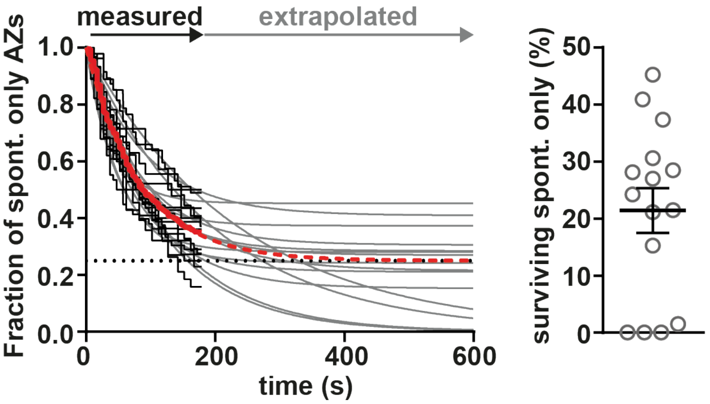

**Figure 2 – figure supplement 1. The majority of BRP-positive AZs are AP-responsive, but a minority is dedicated to spontaneous transmission.** Left: Cell-wise (N =15 animals) survival analysis: AZs found spontaneously active during the first recording episode (1 in Fig. 2A) are tested for their "survival" as "spontaneous only" AZs in the second recording episode (2 in Fig. 2A) where 36 APs are administered at 0.2 Hz. Once a previously spontaneously active AZ responds to an AP, this is considered the "death" of a "spontaneous only" AZ. A fraction of 1 represents all AZs spontaneously active before AP application. Individual black lines indicate the behavior each of the investigated animals/NMJs, the grey lines are fits with a mono exponential decay function that includes a plateau value (see methods for details). The Red line is the fit to the average behavior of all investigated NMJs (N = 15 animals). Right: Fraction of exclusively spontaneously active AZs in relation to all spontaneously active AZs (plateau values from the graph depicted on the left). Circles represent the individual plateau values (fraction of AZs dedicated to spontaneous transmission only) in each of the 15 animals, the vertical line and error bars indicate the mean and SEM. A model with plateau was preferred based on Akaike's Information criterion (see methods for details).

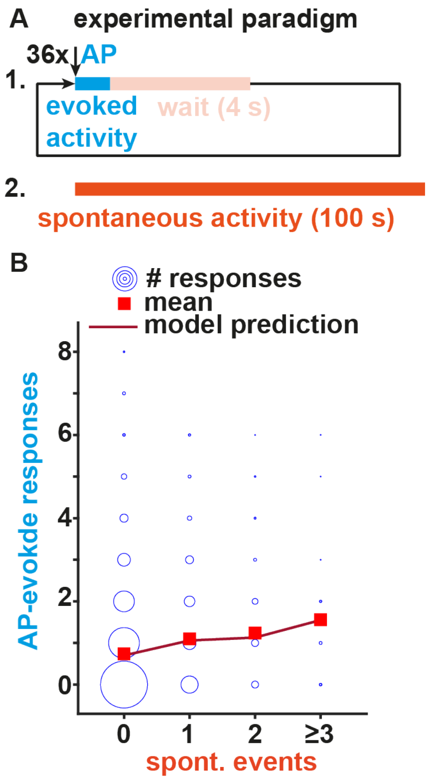

**Figure 2 – figure supplement 2. Inverse experiment where AP-evoked activity is read out before spontaneous activity.** (**A**) Experimental paradigm. 36 APs are elicited at 0.2 Hz (blue) before spontaneous events (orange) are recorded for 100 s in isolation. (**B**) Analysis of the relation between the observed AP-evoked and spontaneous transmission events at individual BRP-positive AZs from 22 animals. Events from all imaged AZs in the 22 animals are pooled. The size of the circles relates to the number of observations (between 1 and 1493). Red squares indicate mean number of evoked responses, the line indicates the model prediction (see methods). Number of BRP-positive AZs investigated: n(AZs)=3194, number of animals in as a function of spontaneous activity: N(0)=22, N(1)=21, N(2)=19, N(≥3)=9).

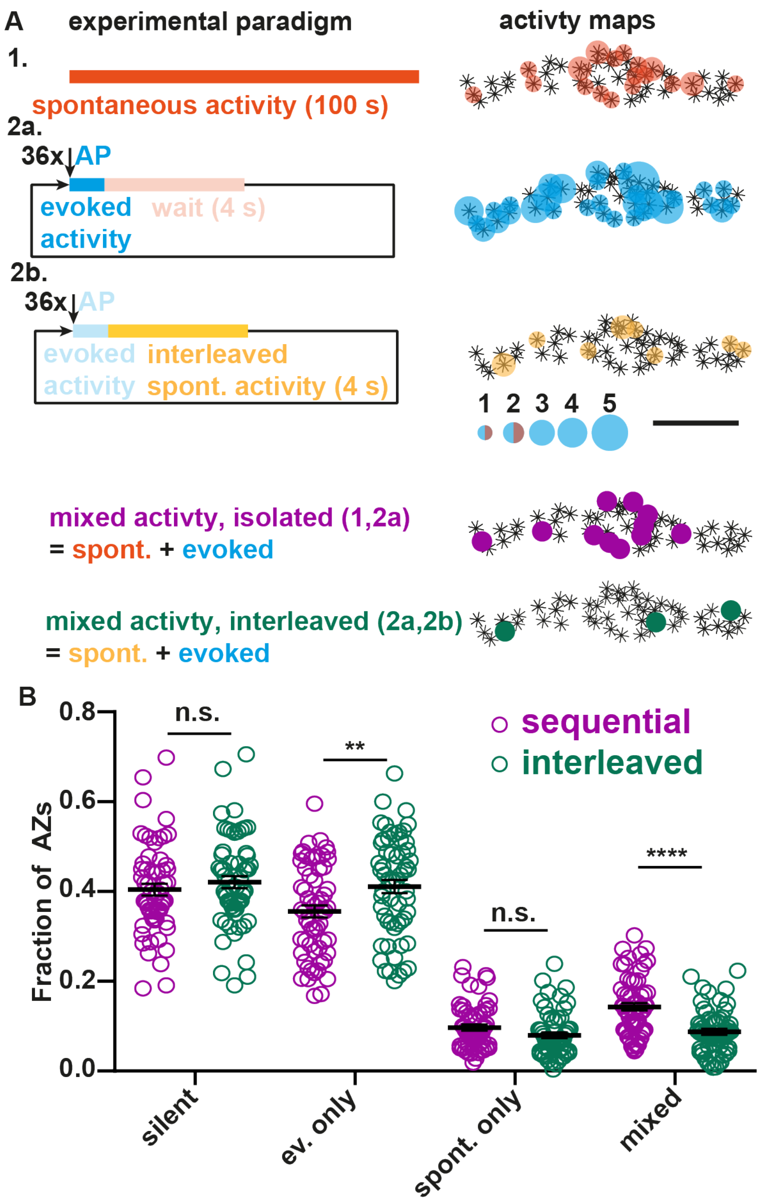

**Figure 2 – figure supplement 3. Spontaneous activity at BRP-positive AZs is reduced when sampled in-between AP stimuli.** (**A**) Left: Two experimental designs to assess spontaneous activity, either in isolation by first sampling spontaneous and then AP-evoked activity (1&2a) or interleaved between the AP-stimuli (2a&2b). Right: Activity maps for isolated spontaneous events (from episode 1), AP-evoked events (from episode 2a) and interleaved spontaneous events (from episode 2b). The same AP-evoked activity (from episode 2a) is used for comparison with the isolated (1) and interleaved (2b) spontaneous activity. (**B**) Animal-wise quantification (N = 59 animals) of fractions of AZs active in the four activity categories (no activity/silent, only AP-evoked activity observed, only spontaneous activity observed, or both activities observed) either measurement sequentially in isolation (purple) or interleaved (green). Lines/Error bars indicate mean and SEM. * <0.05; ** $p<0.01$; *** $p<0.001$; **** $p<0.0001$; n.s. not significant in a two-tailed Student's t-test.

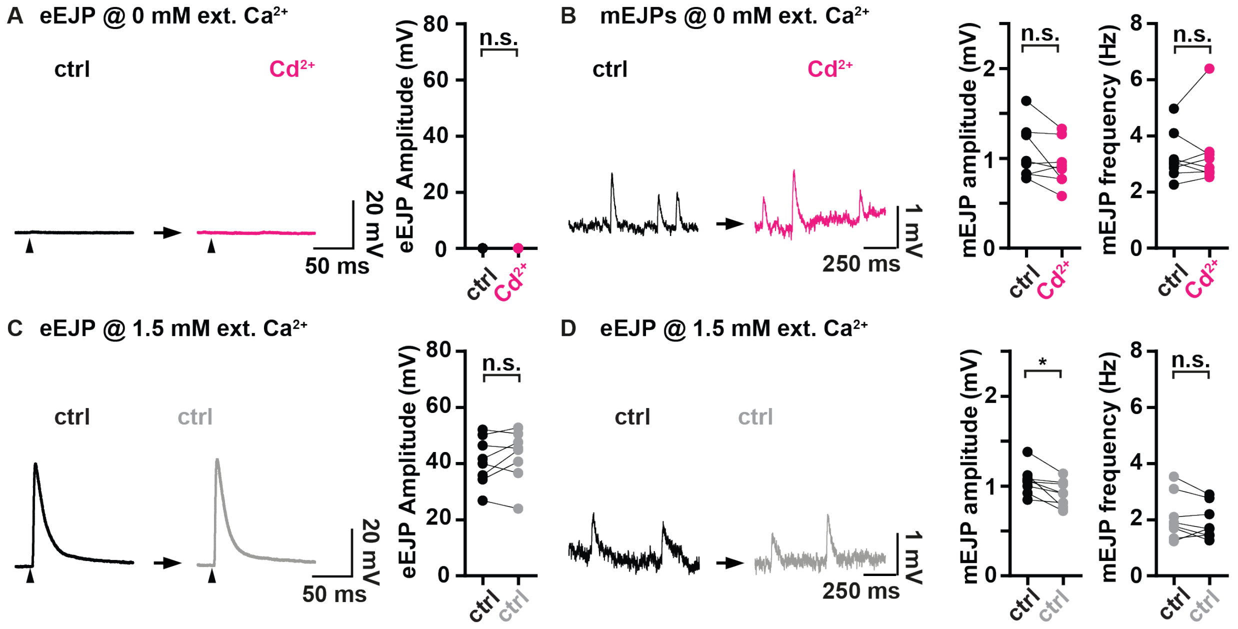

**Figure 3 – figure supplement 1. Application of $Cd^{2+}$ in the absence of extracellular $Ca^{2+}$ has no effects on mEJPs and their frequency is not reduced by the recording paradigm itself.** (**A&B**) Assessment of the consequences of $Cd^{2+}$-application (300 µM, black→magenta) on AP-evoked eEJPs (A) and on spontaneous mEJPs (B) in paired current clamp recordings of muscle 4 NMJs in the absence of extracellular $Ca^{2+}$ (no $Ca^{2+}$ was added to the extracellular solution and 2 mM EGTA was added to buffer any free remaining $Ca^{2+}$). (**A**) Left: Representative eEJP before (black) and after (magenta) application of 300 µM $CdCl_2$. Right: Animal-wise quantification of eEJP amplitudes before (black) and after (magenta) $Cd^{2+}$ application. (**B**) Left: Representative mEJC traces before (black) and after (magenta) $Cd^{2+}$ application. Right: Cell-wise quantification of mEJC amplitudes and -frequencies. (**C&D**) Assessment of the consequences of the experimental paradigm (no treatment, black→grey) on AP-evoked eEJPs (C) and on spontaneous mEJPs (D) in paired current clamp recordings of muscle 4 NMJs in the presence of 1.5 mM $Ca^{2+}$ in the extracellular solution (same condition as in Fig. 3C&D). (**C**) Left: Representative eEJP before (black) and after (grey) application of a mock solution. Right: Animal-wise quantification of eEJP amplitudes before (black) and after (grey) application of a mock solution. (**D**) Left: Representative mEJC traces before (black) and after (grey) application of a mock solution. Right: Cell-wise quantification of mEJC amplitudes and -frequencies. Arrowheads indicate time of stimulation. Number of animals in (A&B and C&D): N=8. n.s. not significant; * $p<0.05$; *** $p<0.001$; **** $p<0.0001$. Paired parametric t-test for comparisons in (A&B and C&D).

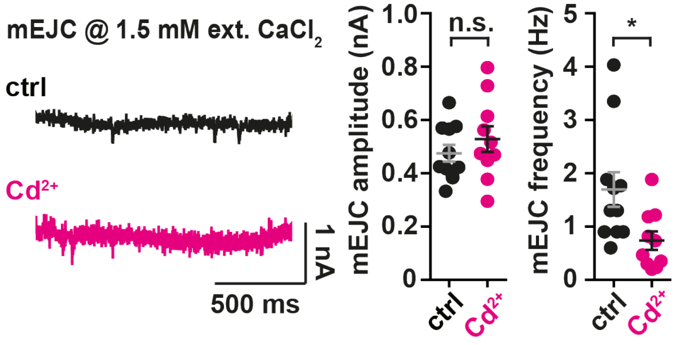

**Figure 3 – figure supplement 2. Investigation of spontaneous transmission upon application of the voltage-gated calcium channel blocker $Cd^{2+}$ in voltage clamp recordings.** Left: Representative mEJC traces of ctrl cells (black) or cells treated with 740.7 µM $Cd^{2+}$ (magenta) recorded in the presence of 1.5 mM extracellular $Ca^{2+}$. Right: Cell-wise quantification of mEJC amplitudes and mEJC frequencies. Number of animals: N(ctrl) = 11, N(Cd2+) = 10. Horizontal bars indicate mean, error bars SEM. n.s. not significant; * $p<0.05$. Two-tailed parametric Student's t-test.

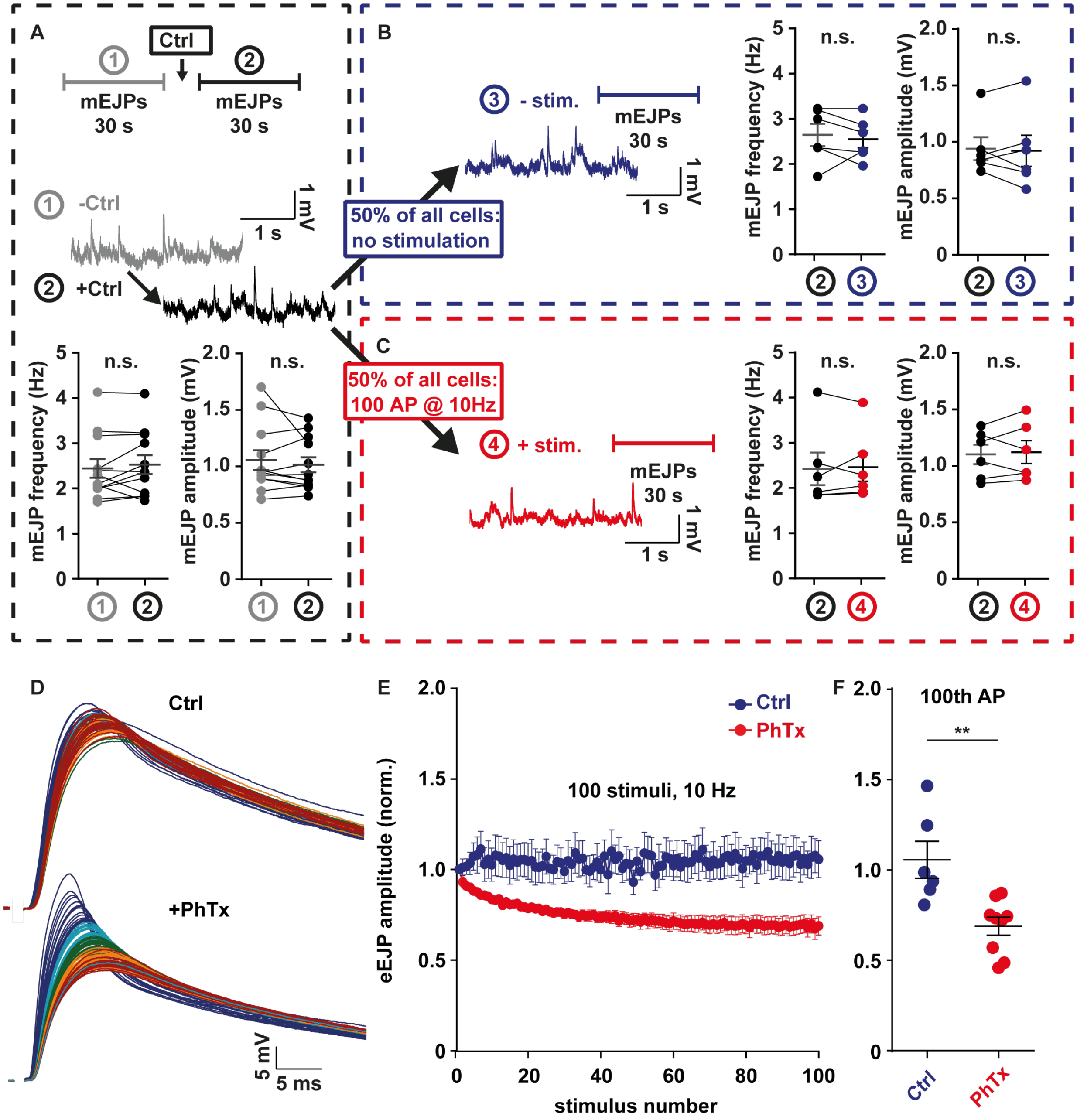

**Figure 4 – figure supplement 1. (A-C)** Analogous to Figure 4, testing whether AP stimulation alone affects spontaneous neurotransmission **D-F**; (**A**) mEJP frequency and amplitude quantification before (grey) and after (black) control treatment (N = 12 animals) (**B**) mEJP frequency and amplitude quantification after control treatment and without stimulation (N = 6 animals) (**C**) mEJP frequency and amplitude quantification after stimulation (10 s wait; N = 6 animals). (**D**) Influence of stimulation on eEJP amplitude over 100 stimuli applied at 10 Hz; top: control conditions, only minor deterioration of amplitudes over 100 APs; bottom: +PhTx, marked decrease of amplitudes over 100 APs. Representative traces shown from 1st eEJP amplitude (blue) to 100th eEJP amplitude (red) (**E**) Sequential quantification of eEJP amplitudes over 100 stimuli applied at 10 Hz in either control (ctrl, blue) or +PhTx (red, 4 µM PhTx) treatment. (**F**) Quantification of 100th eEJP amplitudes in ctrl (blue, N=6 animals) or PhTx conditions (red, N=9 animals). **p<0.01; n.s. not significant. Error bars indicate SEM.

### Supplementary data and statistics

| Figure | Panel | Group | Measure | Mean | SEM | Significance level/alpha | Test type | N (animals) |
|---|---|---|---|---|---|---|---|---|
| **Fig. 1 – fs 1H** | B | 0.4 mM $[Ca^{2+2+}]_{ext}$ | mean GCaMP5G spontaneous event amplitude | 873.390 | 42.670 | | | 9 |
| | | 0.75 mM $[Ca^{2+2+}]_{ext}$ | | 859.783 | 49.161 | | | |
| | | 1.5 mM $[Ca^{2+2+}]_{ext}$ | | 936.153 | 60.632 | | | |
| | | 3 mM $[Ca^{2+2+}]_{ext}$ | | 1084.005 | 62.637 | | | |
| | | 6 mM $[Ca^{2+2+}]_{ext}$ | | 1152.955 | 80.846 | | | |
| | | 12 mM $[Ca^{2+2+}]_{ext}$ | | 1270.339 | 87.833 | | | |

Table 1 – Data relating to Figure 1 – figure supplement 1

| Figure | Measure | Mean | SEM |
|---|---|---|---|
| **Fig. 2 – fs 1** | mean of exp. fit plateaus | 0.2498 | 0.001966 |
| **Fig. 2 – fs 1** | percentage of surviving spont. only Azs | 0.2142 | 0.03192 |

Table 2 – Data relating to Figure 2 – figure supplement 1

| Figure | Panel | Measure | Test type | ANOVA summary | | | Mutliple comparisons | Mean 1 | Mean 1 | Mean Diff. | SE of Diff | N1 | N2 | Adjusted P Value |
|---|---|---|---|---|---|---|---|---|---|---|---|---|---|---|
| **2** | B | Fraction of AP-responsive AZs | Anova | F | 11.9 | | 0 vs. 1 | 0 vs. 1 | 0.467 | -0.1146 | 0.04072 | 59 | 59 | 0.0273 |
| | | | | P value | <0.0001 | | 0 vs. 2 | 0 vs. 2 | 0.467 | -0.1752 | 0.0409 | 59 | 58 | 0.0002 |
| | | | | P value summary | **** | | 0 vs. >2 | 0 vs. >2 | 0.467 | -0.2789 | 0.05016 | 59 | 29 | <0.0001 |
| | | | | Significant diff. Among means (P > 0.05) | Yes | | 1 vs. 2 | 1 vs. 2 | 0.5817 | - 0.06059 | 0.0409 | 59 | 58 | 0.4506 |
| | | | | R square | 0.1518 | | 1 vs. >2 | 1 vs. >2 | 0.5817 | -0.1643 | 0.05016 | 59 | 29 | 0.0068 |
| | | | | Number of values | 205 | | 2 vs. >2 | 2 vs. >2 | 0.6423 | -0.1037 | 0.0503 | 58 | 29 | 0.1693 |
| | | | | Number of treatments | 4 | | | | | | | | | |

Table 3 – Data relating to Figure 2B

| counts | 0 spont | 1 spont | 2 spont | >2 spont |
|---|---|---|---|---|
| **0 ev** | 3966 | 696 | 144 | 14 |
| **1 ev** | 1943 | 490 | 122 | 19 |
| **2 ev** | 890 | 278 | 66 | 7 |
| **3 ev** | 401 | 148 | 33 | 7 |
| **4 ev** | 180 | 55 | 26 | 6 |
| **5 ev** | 78 | 30 | 12 | 2 |
| **6 ev** | 24 | 21 | 3 | 1 |
| **7 ev** | 8 | 2 | 1 | 1 |
| **8 ev** | 2 | 1 | 0 | 0 |
| **Mean evoked activity** | 0.83 | 1.17 | 1.33 | 1.77 |
| **Fitted mean evoked activity** | 0.78 | 1.1 | 1.27 | 1.78 |

Table 4 – Data relating to Figure 2C

| counts | 0 spont | 1 spont | 2 spont | >2 spont |
|---|---|---|---|---|
| **0 ev** | 1493 | 199 | 33 | 5 |
| **1 ev** | 643 | 116 | 34 | 6 |
| **2 ev** | 293 | 76 | 23 | 4 |
| **3 ev** | 110 | 43 | 6 | 1 |
| **4 ev** | 45 | 13 | 3 | 0 |
| **5 ev** | 20 | 6 | 2 | 1 |
| **6 ev** | 5 | 4 | 1 | 1 |
| **7 ev** | 6 | 0 | 0 | 0 |
| **8 ev** | 2 | 0 | 0 | 0 |
| **Mean evoked activity** | 0.74 | 1.10 | 1.24 | 1.56 |
| **Fitted mean evoked activity** | 0.7 | 1.06 | 1.13 | 1.55 |

Table 5 – Data relating to Figure 2 – figure supplement 3

| Figure | Panel | Measure | Group | Mean | SEM | P value | Test type | N (animals) | t | df | F |
|---|---|---|---|---|---|---|---|---|---|---|---|
| **Fig. 2 – fs 3** | B | fraction of all Azs | silent, sequential | 0.4042 | 0.01329 | 0.3711 | Unpaired t test | 59 | 0.8979 | 116 | 1.016 |
| | | | silent, interleaved | 0.4211 | 0.01340 | | | | | | |
| | | | ev only, sequential | 0.3556 | 0.01345 | 0.0061 | | | 2.794 | 116 | 1.193 |
| | | | ev only, interleaved | 0.4113 | 0.01469 | | | | | | |
| | | | sp only, sequential | 0.09686 | 0.006667 | 0.0704 | | | 1.826 | 116 | 1.066 |
| | | | sp only, interleaved | 0.07991 | 0.006456 | | | | | | |
| | | | mixed, sequential | 0.1433 | 0.008586 | <0.0001 | | | 5.169 | 116 | 1.742 |
| | | | mixed, interleaved | 0.8765 | 0.006504 | | | | | | |

Table 6 – Data relating to Figure 2 – figure supplement 3

| Figure | Measure | Group | Mean | SEM | Significance level/alpha | Test type | Comment | N (animals) |
|---|---|---|---|---|---|---|---|---|
| **Fig. 3A** | average GCaMP fluorescence amplitude | ctrl | 371.7 | 20.18 | 0.5573 | unpaired non-parametric t-test | | 18 |
| | | TTX | 353.5 | 20.45 | | | | |
| | avg. spont. event frequency (Hz/AZ) | ctrl | 0.00217 | 0.0002246 | 0.7123 | unpaired non-parametric t-test | | |
| | | TTX | 0.002162 | 0.0003297 | | | | |
| **Fig. 3B** | average GCaMP fluorescence amplitude | ctrl | 409.4 | 35.19 | 0.0668 | unpaired non-parametric t-test | | 11 |
| | | Cd | 330.9 | 18.01 | | | 2 animals showed no events | 9 |
| | avg. spont. event frequency (Hz/AZ) | ctrl | 0.00221 | 0.000354 | < 0.0001 | unpaired non-parametric t-test | | 11 |
| | | Cd | 0.0002248 | 0.00007827 | | | | 11 |

Table 7 – Data relating to Figure 3A,B

| Figure | Measure | Group | Mean | SEM | Significance level/alpha | Test type | N (animals) |
|---|---|---|---|---|---|---|---|
| **Fig. 3C** | eEJP amplitude (mV), 1.5 mM $[Ca^{2+}]_{ext.}$ | ctrl before | 51.73 | 2.242 | <0.0001 | paired parametric t-test | 8 |
| | | $CdCl_2$ | 0 | 0 | | | |
| **Fig. 3D** | mEJP amplitude (mV), 1.5 mM $[Ca^{2+}]_{ext.}$ | ctrl before | 1.046 | 0.07287 | <0.0001 | paired parametric t-test | 8 |
| | | $CdCl_2$ | 0.7458 | 0.05095 | | | |
| | mEJP frequency(Hz), 1.5 mM $[Ca^{2+}]_{ext.}$ | ctrl before | 3.013 | 0.5100 | 0.0068 | paired parametric t-test | 8 |
| | | $CdCl_2$ | 2.250 | 0.4348 | | | |

Table 8 – Data relating to Figure 3C,D

| Figure | Panel | Measure | Group | Mean | SEM | Significance level/alpha | Test type | Comment | N (animals) |
|---|---|---|---|---|---|---|---|---|---|
| **Fig. 3 – fs 1** | A,C | eEJP amplitude (mV), 0 mM $[Ca^{2+}]_{ext.}$ | ctrl before | 0 | 0 | | | no test could be performed since all values were 0 | 8 |
| | | | ctrl after | 0 | 0 | | | | |
| | | | ctrl before | 0 | 0 | | | | |
| | | | $CdCl_2$ | 0 | 0 | | | | |
| **Fig. 3 – fs 1** | C | mEJP amplitude (mV), 1.5 mM $[Ca^{2+}]_{ext.}$ | ctrl before | 1.057 | 0.05586 | 0.0121 | paired parametric t-test | | 8 |
| | | | ctrl after | 0.9178 | 0.05121 | | | | |
| | B,D | mEJP amplitude (mV), 0 mM $[Ca^{2+}]_{ext.}$ | ctrl before | 0.9713 | 0.1602 | 0.1467 | paired parametric t-test | | 8 |
| | | | ctrl after | 0.8994 | 0.1329 | | | | |
| | C | mEJP frequency(Hz), 1.5 mM $[Ca^{2+}]_{ext.}$ | ctrl before | 2.083 | 0.2898 | 0.2543 | paired parametric t-test | | 8 |
| | | | ctrl after | 1.925 | 0.2211 | | | | |
| | D | mEJP frequency (Hz), 0 mM $[Ca^{2+}]_{ext.}$ | ctrl before | 2.421 | 0.1776 | 0.4166 | paired parametric t-test | | 8 |
| | | | ctrl after | 2.508 | 0.2175 | | | | |

Table 9 – Data relating to Figure 3 – figure supplement 1

| Figure | Measure | Group | Mean | SEM | Significance level/alpha | Test type | Comment | N (animals) |
|---|---|---|---|---|---|---|---|---|
| **Fig. 3 fs 2** | mEJC amplitude (nA) | ctrl | 0.4754 | 0.03126 | 0.3618 | unpaired parametric t-test | | 11 |
| | | Cd | 0.5282 | 0.04835 | | | | 10 |
| | mEJC frequency (Hz) | ctrl | 1.695 | 0.3255 | 0.0209 | unpaired parametric t-test | | 11 |
| | | Cd | 0.7383 | 0.1734 | | | | 10 |

Table 10 – Data relating to Figure 3 – figure supplement 2

| Figure | Panel | Measure | Group | Mean | SEM | Significance evel/alpha | Test type | Comment | N (animals) |
|---|---|---|---|---|---|---|---|---|---|
| **Fig. 4** | D | mEJP frequency (Hz) | before PhTx | 2.417 | 0.1329 | 0.0435 | paired parametric t-test | | 18 |
| | | mEJP frequency (Hz) | after PhTx | 2.226 | 0.1917 | | | | 18 |
| **Fig. 4** | D | mEJP amplitude (mV) | before PhTx | 0.8902 | 0.03051 | < 0.0001 | paired parametric t-test | | 18 |
| | | mEJP amplitude (mV) | after PhTx | 0.5702 | 0.02297 | | | | 18 |
| **Fig. 4** | E | mEJP frequency (Hz) | after PhTx | 2.163 | 0.2705 | 0.8962 | paired parametric t-test | | 9 |
| | | mEJP frequency (Hz) | no stim | 2.144 | 0.2429 | | | | 9 |
| **Fig. 4** | E | mEJP amplitude (mV) | after PhTx | 0.5261 | 0.03291 | 0.8328 | paired parametric t-test | | 9 |
| | | mEJP amplitude (mV) | no stim | 0.5218 | 0.02073 | | | | 9 |
| **Fig. 4** | F | mEJP frequency (Hz) | after PhTx | 2.289 | 0.2863 | 0.0027 | paired parametric t-test | | 9 |
| | | mEJP frequency (Hz) | stim | 1.726 | 0.179 | | | | 9 |
| **Fig. 4** | F | mEJP amplitude (mV) | after PhTx | 0.6144 | 0.02594 | 0.2031 / 0.1371 | Wilcoxon matched-pairs signed rank test / paired parametric t-test | Group “after PhTx” matched with “stimulation” group failed D’Agostino & Pearson omnibus normality test | 9 |
| | | mEJP amplitude (mV) | stim | 0.5675 | 0.04475 | | | | 9 |

Table 11 – Data relating to Figure 4

| Figure | Panel | Measure | Group | Mean | SEM | Significance evel/alpha | Test type | Comment | N (animals) |
|---|---|---|---|---|---|---|---|---|---|
| **Fig. 4 – fs 1** | A | mEJP frequency (Hz) | before ctrl | 2.444 | 0.2086 | 0.3491 | paired nonparametric t-test | | 12 |
| | | mEJP frequency (Hz) | after ctrl | 2.528 | 0.2091 | | | | 12 |
| **Fig. 4 – fs 1** | A | mEJP amplitude (mV) | before ctrl | 1.057 | 0.08722 | 0.5186 | paired nonparametric t-test | | 12 |
| | | mEJP amplitude (mV) | after ctrl | 1.014 | 0.06722 | | | | 12 |
| **Fig. 4 – fs 1** | B | mEJP frequency (Hz) | after ctrl | 2.650 | 0.2426 | 0.6250 | paired nonparametric t-test | | 6 |
| | | mEJP frequency (Hz) | after ctrl, -stim | 2.556 | 0.1891 | | | | 6 |
| **Fig. 4 – fs 1** | B | mEJP amplitude (mV) | after ctrl | 0.9385 | 0.1011 | 0.8438 | paired nonparametric t-test | | 6 |
| | | mEJP amplitude (mV) | after ctrl, -stim | 0.9202 | 0.1380 | | | | 6 |
| **Fig. 4 – fs 1** | C | mEJP frequency (Hz) | after ctrl | 2.406 | 0.3572 | 0.8438 | paired nonparametric t-test | | 6 |
| | | mEJP frequency (Hz) | after ctrl, +stim | 2.444 | 0.3129 | | | | 6 |
| **Fig. 4 – fs 1** | C | mEJP amplitude (mV) | after ctrl | 1.090 | 0.08576 | 0.6875 | paired nonparametric t-test | | 6 |
| | | mEJP amplitude (mV) | after ctrl, +stim | 1.109 | 0.1024 | | | | 6 |
| **Fig. 4 – fs 1** | F | norm. eEJP amplitude | ctrl | 1.057 | 0.1020 | 0.0016/0.0033 | Mann-Whitney U test/unpaired parametric t-test | | 6 |
| | | | PhTx | 0.6888 | 0.04986 | | | | 9 |

Table 12 – Data relating to Figure 4 – figure supplement 1
